# Sox2 controls neural stem cell self-renewal through a Fos-centered gene regulatory network

**DOI:** 10.1101/2020.03.17.995621

**Authors:** Miriam Pagin, Mattias Pernebrink, Simone Giubbolini, Cristiana Barone, Gaia Sambruni, Yanfen Zhu, Matteo Chiara, Sergio Ottolenghi, Giulio Pavesi, Chia-Lin Wei, Claudio Cantù, Silvia K. Nicolis

## Abstract

The Sox2 transcription factor is necessary for the long-term self-renewal of neural stem cells (NSC). Its mechanism of action is still poorly defined. To identify molecules regulated by Sox2, and acting in mouse NSC maintenance, we transduced, into Sox2-deleted NSC, genes whose expression is strongly downregulated following Sox2 loss (Fos, Jun, Egr2), individually or in combination. Fos alone rescued long-term proliferation, as shown by in vitro cell growth and clonal analysis. Further, pharmacological inhibition of the FOS/JUN AP1 complex binding to its targets, decreased cell proliferation and expression of the putative target Suppressor of cytokine signaling 3 (Socs3). Additionally, Fos requirement for efficient long-term proliferation was demonstrated by the reduction of NSC clones capable of long-term expansion following CRISPR/Cas9-mediated Fos inactivation. Previous work showed that the Socs3 gene is strongly downregulated following Sox2 deletion, and its reexpression by lentiviral transduction rescues long-term NSC proliferation. Fos appears to be an upstream regulator of Socs3, possibly together with Jun and Egr2; indeed, Sox2 reexpression in Sox2-deleted NSC progressively activates both Fos and Socs3 expression; in turn, Fos transduction activates Socs3 expression. Based on available SOX2 ChIPseq and ChIA-PET data, we propose a model whereby Sox2 is a direct activator of both Socs3 and Fos, as well as possibly Jun and Egr2; further, we provide direct evidence for FOS and JUN binding on *Socs3* promoter, suggesting direct transcriptional regulation. These results provide the basis for developing a model of a network of interactions, regulating critical effectors of NSC proliferation and long-term maintenance.

**Significance statement:** Proliferation and maintenance of NSC are essential during normal brain development, and, postnatally, for the maintenance of hippocampal function and memory until advanced age. Little is known about the molecular mechanisms that maintain the critical aspects of NSC biology (quiescence and proliferation) in postnatal age. Our work provides a methodology, transduction of genes deregulated following Sox2 deletion, that allows to test many candidate genes for their ability to sustain NSC proliferation. In principle, this may have interesting implications for identifying targets for pharmacological manipulations.

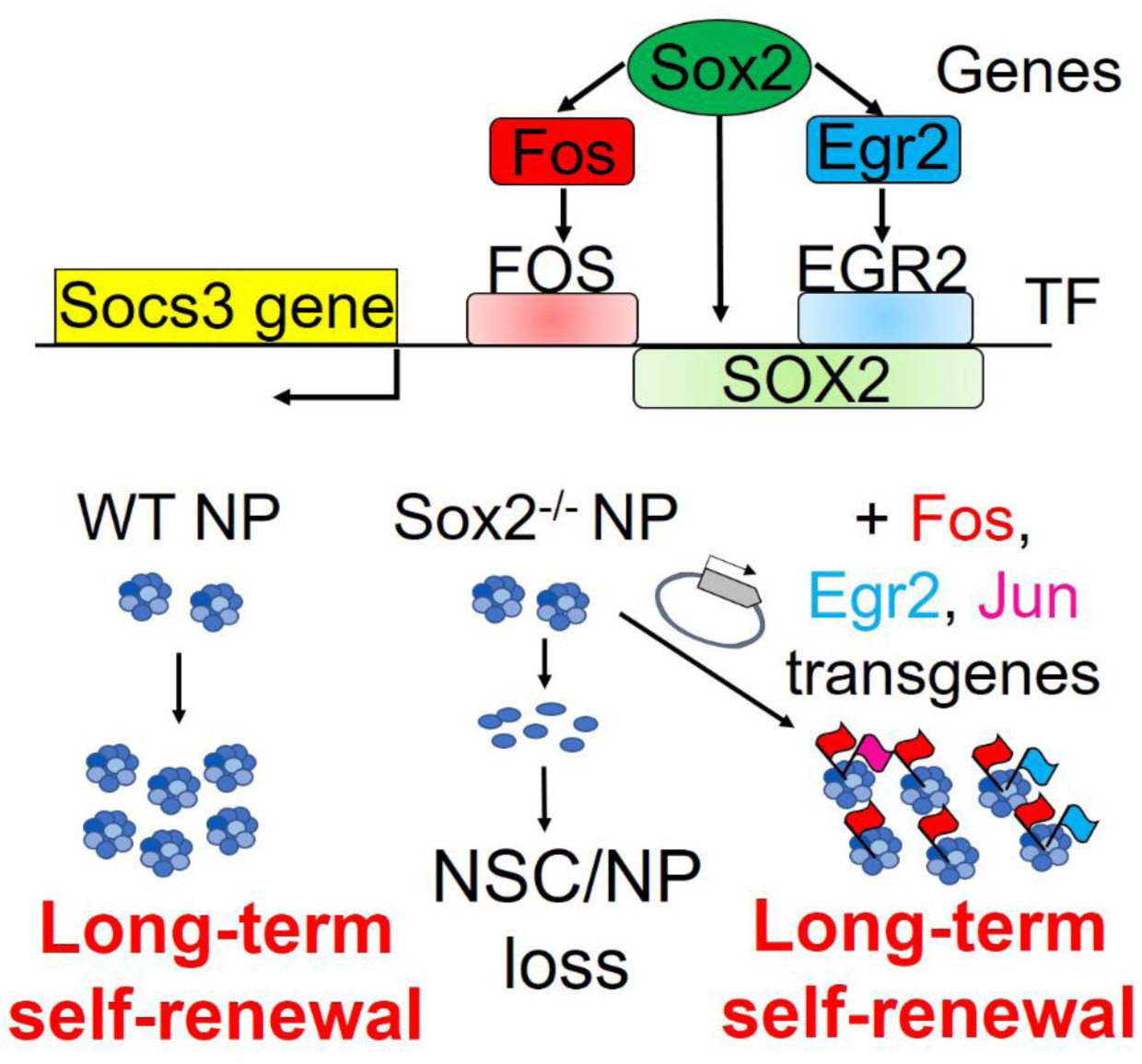

## Introduction

The transcription factor Sox2 is critically important in the development of the brain. In humans, Sox2 mutations cause brain abnormalities (hippocampal dysplasia, learning disabilities, epilepsy, motor control problems, eye and vision defects) [1-4]. Sox2 is expressed in neural stem and progenitor cells; its deletion during mouse embryogenesis causes prenatal or early postnatal loss of NSC in the hippocampus, and its postnatal loss decreases hippocampal neural stem/progenitor cell proliferation [5]. Molecular mechanisms of Sox2 roles in supporting NSC maintenance have been widely investigated [3,6-12]. In vitro experiments on NSC showed that Sox2 loss causes progressive exhaustion of NSC, in contrast to the long-term proliferation of control wild type cells [5]; differentiation defects of progenitor cells have also been reported in Sox2-mutant cells by [13] and [8]. The loss of self-renewal in Sox2-mutant cells grown in vitro is reversible; in fact, in Sox2-deleted cells, re-expression of Socs3, the gene most downregulated following Sox2 loss, rescued long-term self-renewal of mutant cells [14]. These findings point to the possibility of identifying the network of regulatory processes acting downstream to Sox2 in allowing NSC proliferation and preserving their functional integrity. To this end, we expressed, in Sox2-deleted NSC, some of the most downregulated genes identified by RNAseq comparison of Sox2-deleted and wild type NSC. We show that transduction of the Fos transcription factor increases Socs3 levels and efficiently rescues long-term proliferation of Sox2-deleted NSC. Pharmacological inhibition of FOS transcriptional activity in wild-type NSC reduces their proliferation, together with the expression of Socs3 mRNA; further, mutation of Fos decreases long-term self-renewal of NSC. Moreover, mining of our previous NSC genomic analyses of SOX2 ChIPseq and RNApolII ChIA-PET, detecting long-range promoter-enhancer chromatin interactions [14], further shows that the genes encoding Fos and its partner Jun, and Egr2 (a regulator of Socs3), are all bound by SOX2. Finally, the Socs3 promoter is bound, in NSC chromatin, by SOX2, JUN and FOS. These experiments define a gene regulatory network (GRN) of interactions contributing to Sox2-dependent long-term NSC self-renewal.

## Materials and Methods

### Primary ex-vivo neural stem/progenitor cell cultures

Brain-derived NSC cultures were obtained from dissected telencephalon of wild-type and Sox2-deleted P0 mice, and grown in fresh medium (FM): DMEM-F12 with Glutamax (GIBCO), supplemented with 1/50 vol/vol B27 (Life Technologies), 1% of Penicillin-Streptomycin (Euroclone) supplemented with EGF (10 ng/ml, Tebu-bio) and bFGF (10 ng/ml, Tebu-bio) as mitogen, essentially as described in [5,14,15].

### Lentiviral constructs

Fos, Jun and Egr2 cDNAs were derived from Addgene plasmids: Fos: “pcDNA3FLAGFos WT”, from John Blenis (#8966); Jun: “FlagJunWTMyc”, from Axel Behrens (#47443); Egr2:”mEgr2/LZRS”, from David Wiest (#27784). cDNAs were amplified by PCR and cut with BamHI, and cloned into a unique BamHI site upstream to the IRES-dNGFR cassette of the pHR SIN BX IR/EMW [16](a gift from A. Ronchi, Milano). The Sox2-expressing lentivirus was previously described [5]; it also expresses the GFP fluorescent marker.

Lentiviral vectors were produced by calcium phosphate transfection into the packaging human embryonic kidney cell line 293T, of the VSV-G plasmid (encoding ENV), CMV R8.74 (packaging) and pRSV-REV (encoding reverse transcriptase)[14,16]. Briefly, after transfection, following replacement with DMEM high glucose (Euroclone), containing 10% Fetal Bovine Serum (Sigma), 1% penicillin-streptomycin (Euroclone), 1% of L-Glutamine (Euroclone), the cell supernatants were collected 24-48 hours after transfection. Lentiviral vectors were titrated on HEK-293T cells by measuring the percentage of eGFP (for the Sox2-transducing vector) or of dNGFR-positive cells (for the Fos, Jun, Egr2-transducing vectors) by Flow Cytometry [16].

### NSC transduction with lentiviral constructs encoding Fos, Jun, Egr2, or Empty Vector

NSC transduction [14]: wild-type and Sox2-deleted neurospheres were grown for 2-3 passages (3-4 days each) in FM supplemented with bFGF (10ng/ml, Tebu-bio) and EGF (10 ng/ml, Tebu-bio), and for one more passage in FM supplemented with EGF only. For passaging, neurospheres were first incubated in 0.25% trypsin (GIBCO) for 5 minutes at 37°C and, subsequently, ovomucoid (Leibovitz’s L15 medium (GIBCO) containing trypsin inhibitor (Sigma), BSA (Sigma) and 40 µg/ml DNaseI (Sigma)) for 5 minutes at 37°C. Spheres were then carefully dissociated mechanically by gently pipetting up and down, centrifuged at 1200 rpm for 4 minutes and cells were resuspended in 1ml of FM, counted and replated at a density of 25,000 cells/1ml/well (9 wells for each cell type) in 24-well plates, in FM with EGF. After 4 hours cells (Sox2-deleted or wild-type) were transduced with appropriate lentiviral vectors (single or in combination), at a multiplicity of infection (MOI) 5.5 for each vector. Cells were incubated overnight at 37°C, then 1ml per well of FM with EGF was added both to transduced and non-transduced (control) cells. After 4 days, cells were dissociated to single cells as above, counted, and seeded at a density of 25,000 cells/well in FM with EGF (routinely, 9 wells /experimental sample) (passage 1). Subsequently, cells were grown for 3.5 days, neurospheres were dissociated and cells counted; an aliquot (25.000 cells/1 ml/well, 9 wells, for Fos, Fos+Egr2, Fos+Jun transduction experiments) was again plated, grown for 3.5 days, counted and again passaged as above, for several additional passages, to obtain a cumulative growth curve (Fig 2A, 2B). The number of cells at the end of each passage, divided by the number of cells plated at the beginning of the passage, measures the cell expansion/passage; by multiplying between them these ratios at subsequent stages of the culture, the overall expansion of the cells is obtained, and plotted as a cumulative growth curve. In addition, at various stages of the culture, 500,000 cells for each sample (from pooled wells) were fixed using PFA 4% and stained as in [16] with an anti-human CD271 (dNGFR) antibody, conjugated with Phycoerythrin (PE) (BioLegend, Cat. No. 345106, RRID: AB_2152647) (dilution 1:200) and analyzed by CytoFLEX (Beckman-Coulter) to determine the percentage of infected cells: 10,000 events were analyzed for each sample.

**Figure 1.**
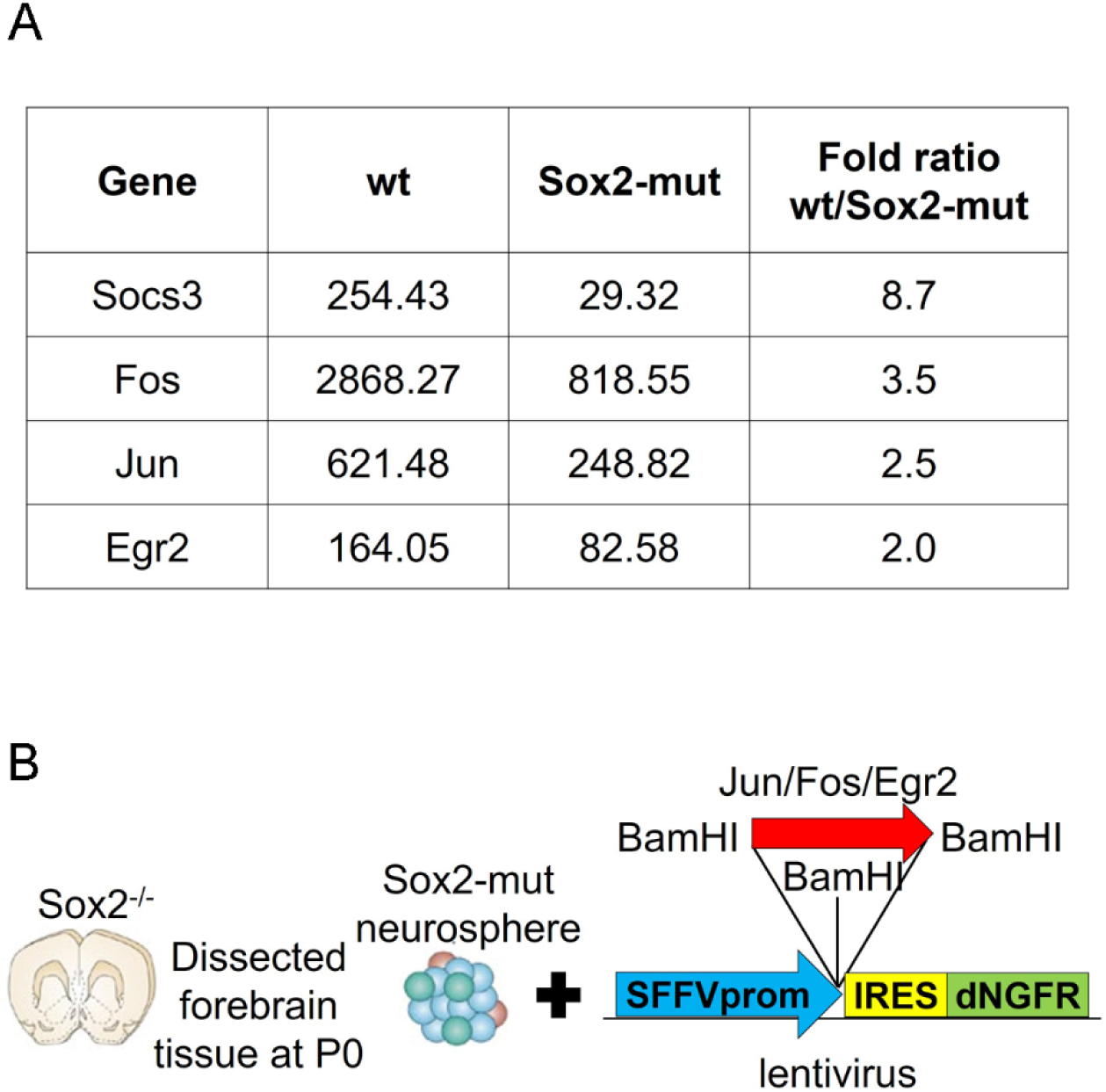
A, Expression levels of selected transcription factor genes among the most downregulated genes in Sox2-deleted NSC (transcripts per million; data from [14]). **B, Scheme of the Sox2 target transduction experiment and of the lentiviral vector**

**Figure 2.**
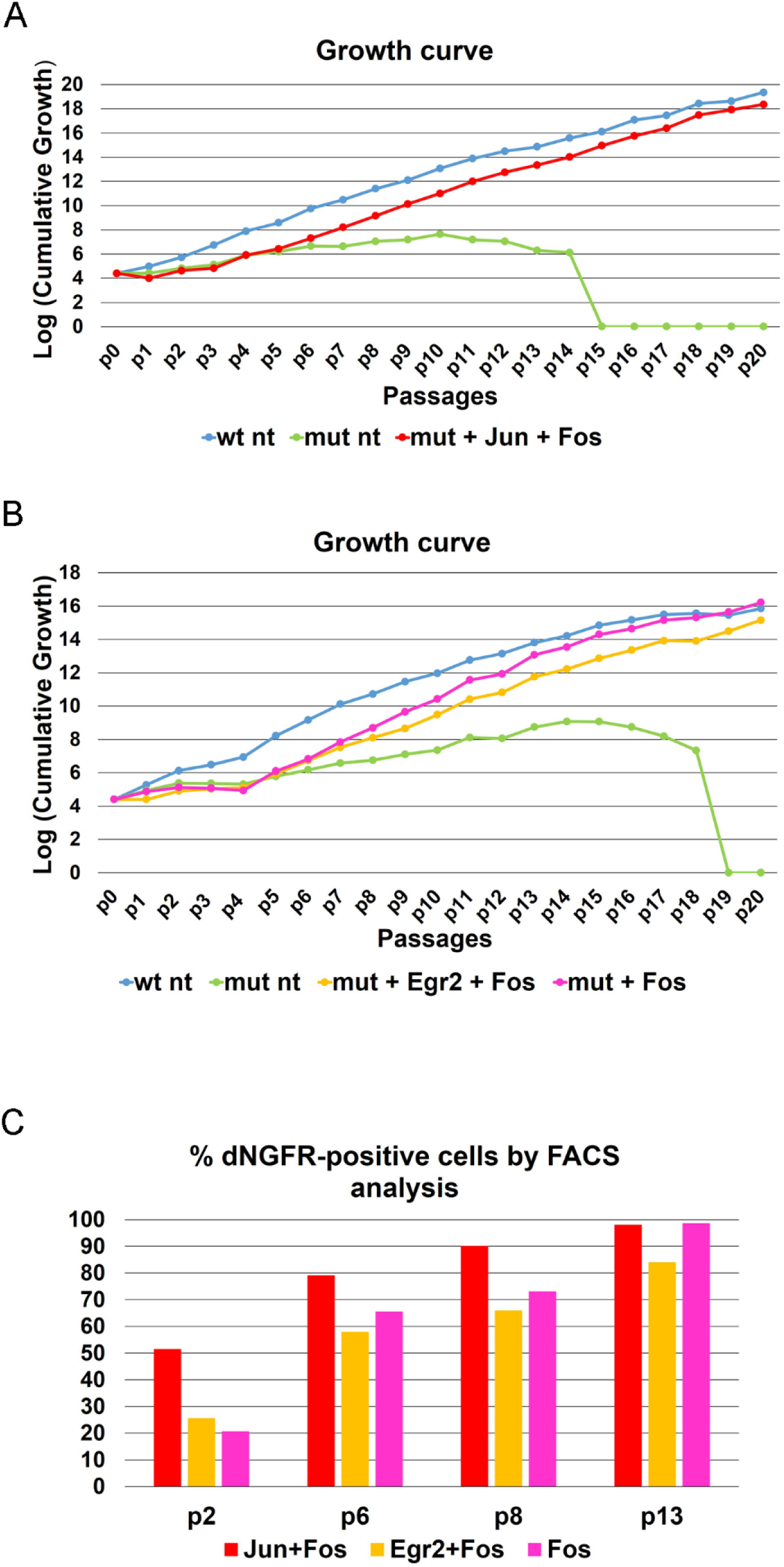
Sox2-mutant cells transduced with Fos+Jun, Fos+Egr2, or Fos alone recover the ability to self-renew long term. **A**,**B**, growth curves of Sox2-deleted cells (green), and of the same cells transduced with the indicated vectors. We performed two independent transductions with Fos alone of Sox2-deleted NSC; the results obtained with the second transduction (not shown) are essentially the same as those depicted here with the first transduction. **C**, Percentage of dNGFR-positive cells by FACS analysis .

### Identification of Fos, Jun, Egr2 lentivirus integrated in clones of lentivirally transduced NSC

Cell populations of NSC transduced with lentiviruses encoding Fos, Jun, Egr2, obtained after 6 or more passages from the transduction, were plated at limiting dilution into 96-well plates (down to a dilution of 7.5 cells/well, which gave either one or no clone in every well). Individual clones (neurospheres) were analyzed by PCR following a short expansion (2-3 passages), DNA was extracted and analyzed by PCR using primers designed to detect the virally transduced, but not the endogenous, Fos, Jun and Egr2 (forward primer is in the viral SFFV promoter driving the cDNA expression; reverse primer is in the cDNA).

Forward primer SFFV: 5’-CTCACAACCCCTCACTCG-3’

Fos Reverse primer: 5’-AGGTCATTGGGGATCTTGCA-3’ (and 5’-GGCTGGGGAATGGTAGTAGG-3’)

Jun Reverse primer: 5’-GGTTCAAGGTCATGCTCTGT-3’

Egr2 Reverse primer: 5’-TGCCCGCACTCACAATATTG-3’

### Transduction of Sox2-deleted NSC with Sox2-encoding lentiviral vector

Sox2-deleted NSCs were dissociated to single cells and seeded at a density of 5×10 ^5^ cells/T25cm^2^ flask/5ml, in FM. After 4 hours NSCs were transduced with a GFP-Sox2-expressing lentivirus [5], at a multiplicity of infection (MOI) 8 and 12 (Suppl. Fig. 3). Cells were incubated overnight at 37°C. Virus was removed by medium change at 24 h: cells were centrifuged at 1000 rpm for 4 minutes and resuspended in FM with EGF. Non-transduced controls were treated equivalently (without virus). At every passage, every 3-4 days, cells were dissociated to single cells as above, counted, and replated in FM with EGF at a density of 20,000 cells/well/1ml, to generate a cumulative growth curve (Suppl. Fig 3). At successive passages, aliquots of Sox2-transduced cells (50,000-500,000 cells, from pooled wells) were fixed using 4% paraformaldehyde (PFA) and analyzed for GFP fluorescence by CytoFLEX (Beckman-Coulter) to determine the percentage of infected cells: 10,000 events were analyzed for each sample.

**Figure 3.**
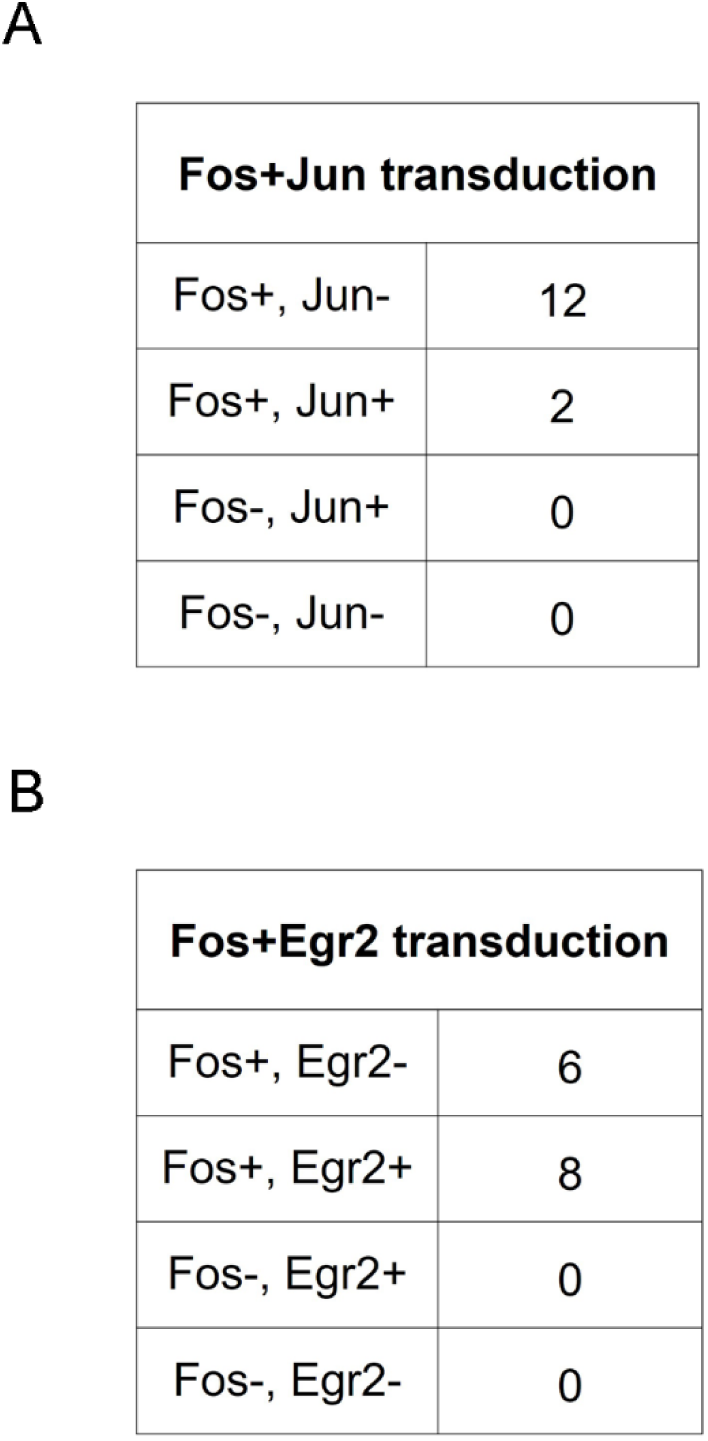
Clones cultured from Sox2-mutant cells rescued with Fos+Jun (A), or Fos+Egr (B), always contain the Fos-expressing lentivirus. Numbers of clones are indicated. The transduction efficiency of the cells by Jun alone, in preliminary experiments, was 18.5%; hence, we would expect that 18.5% of the 14 Fos-positive clones, i.e 2-3 clones, should also be Jun-positive, by chance. For Egr2, the expected frequency of Fos/Jun positive clones is 17.1% of the 14 Fos-positive clones, i.e. 2-3 clones.

### Measure of Fos and Socs3 expression levels in Sox2-deleted cells transduced with Sox2-or Fos-encoding lentiviral vectors

Sox2-deleted NSCs were dissociated to single cells and seeded at a density of 5×10 ^5^ cells/T25cm^2^ flask/5ml, in FM. After 4 hours NSCs were transduced with lentiviruses expressing Sox2 or Fos at a multiplicity of infection (MOI) 10. Cells were incubated overnight at 37°C. Virus was removed by medium change at 24 h: cells were centrifuged at 1000 rpm for 4 minutes and resuspended in FM. Non-transduced controls were treated equivalently (without virus).

At every passage, every 3-4 days, cells were dissociated to single cells as above, counted, and replated in FM with EGF at a density of 800,000 cells/T75cm^2^/12ml.; 50,000-500,000 cells were analyzed by FACS for GFP or dNGFR expression, as described above, to determine the percentage of transduced cells. Additionally, aliquots of cells (100,000-500,000 cells) were lysed in TRIzol (Life Technologies) for total RNA extraction. RNA was purified using Direct-zol RNA Miniprep (Zymo Research), treated with DNase (Zymo Research) for 30 minutes at 37° C and 400ng of RNA were retrotranscribed (High Capacity cDNA Reverse Transcription Kit, Applied Biosystem). About 5μl of a 1:25 dilution (adjusted following normalization by *Hprt* expression) were used for the real time PCR. Negative control reactions (without Reverse Transcriptase) gave no PCR amplification. Real time analysis was performed using ABI Prism 7500 (Applied Biosystems). Samples from each experiment were analyzed in duplicate. Specific PCR product accumulation was monitored by using SsoAdvanced™ Universal SYBR® Green Supermix (Bio-Rad) fluorescent dye in a 12-μl reaction volume. Dissociation curves confirmed the homogeneity of PCR products.

Primers for the quantification of Fos and Socs3 mRNAs are:

Fos Forward primer: 5’-CTGTCCGTCTCTAGTGCCAA-3’

Fos Reverse primer: 5’-TGCTCTACTTTGCCCCTTCT-3’

Socs3 Forward primer: 5’-ACCTTTGACAAGCGGACTCT-3’

Socs3 Reverse primer: 5’-AGGTGCCTGCTCTTGATCAA-3’

HPRT Forward primer: 5’-TCCTCCTCAGACCGGTTT-3’

HPRT Reverse primer: 5’-CCTGGTTCATTCATCGCTAATC-3’

All qRT-PCRs for Fos and Socs3 were performed in parallel with HPRT qRT-PCR, for normalization.

### CRISPR-Cas9 assays

Wild-type neurospheres were dissociated into single cells, as described above, and seeded at a density of 5×10^5^ cells/T25cm^2^ flask/5ml, in FM with EGF. Cells were transduced, after 4 hours from plating, with a previously defined amount of lentiCRISPRV2puro lentivirus expressing the anti-Fos guide RNA or the scramble non-targeting guide RNA. Medium was changed after 3 days and cells were selected in FM with 5μg/ml Puromycin (Sigma-Aldrich, cod. P8833) for 3 days. To remove puromycin, cells were then centrifuged at 1000 rpm for 4 minutes and resuspended in FM with EGF without puromycin and grown for 15 days to allow recovery from the stress of transduction and selection. Neurospheres were then dissociated to single cells; similar numbers of cells treated with scramble guide RNA (Mock) or with anti-Fos guide RNA (Mut) were plated at clonal dilution in individual wells of a 96-well plate in FM with EGF. With 15 cells/well, essentially all wells showed, after 10 days, at least one or more neurospheres; at a nominal limiting dilution of 7.5 cells/well, single clones (neurospheres) appeared only in a proportion of the wells (the “primary clones”). These clones were then further used to determine the proportion of long-term growing clones (see below). Two experiments were performed: 258 and 207 primary clones were obtained from cells treated with scramble guide RNA; 73 and 198 primary clones were detected from cells treated with anti-Fos guide RNA.

Individual primary clones from wells showing a single neurosphere were picked up, disaggregated and replated (“replated clones”, Fig. 5C, 5D) in a single well of a 48-well plate, for further expansion in a 24-well plate and, finally, in a 6-well plate and in flasks. At every passage, the number of clones which did not proliferate was annotated. Clones that continued to grow in flasks or 6-well plates were considered to be “long-term growing clones”.

**Figure 4.**
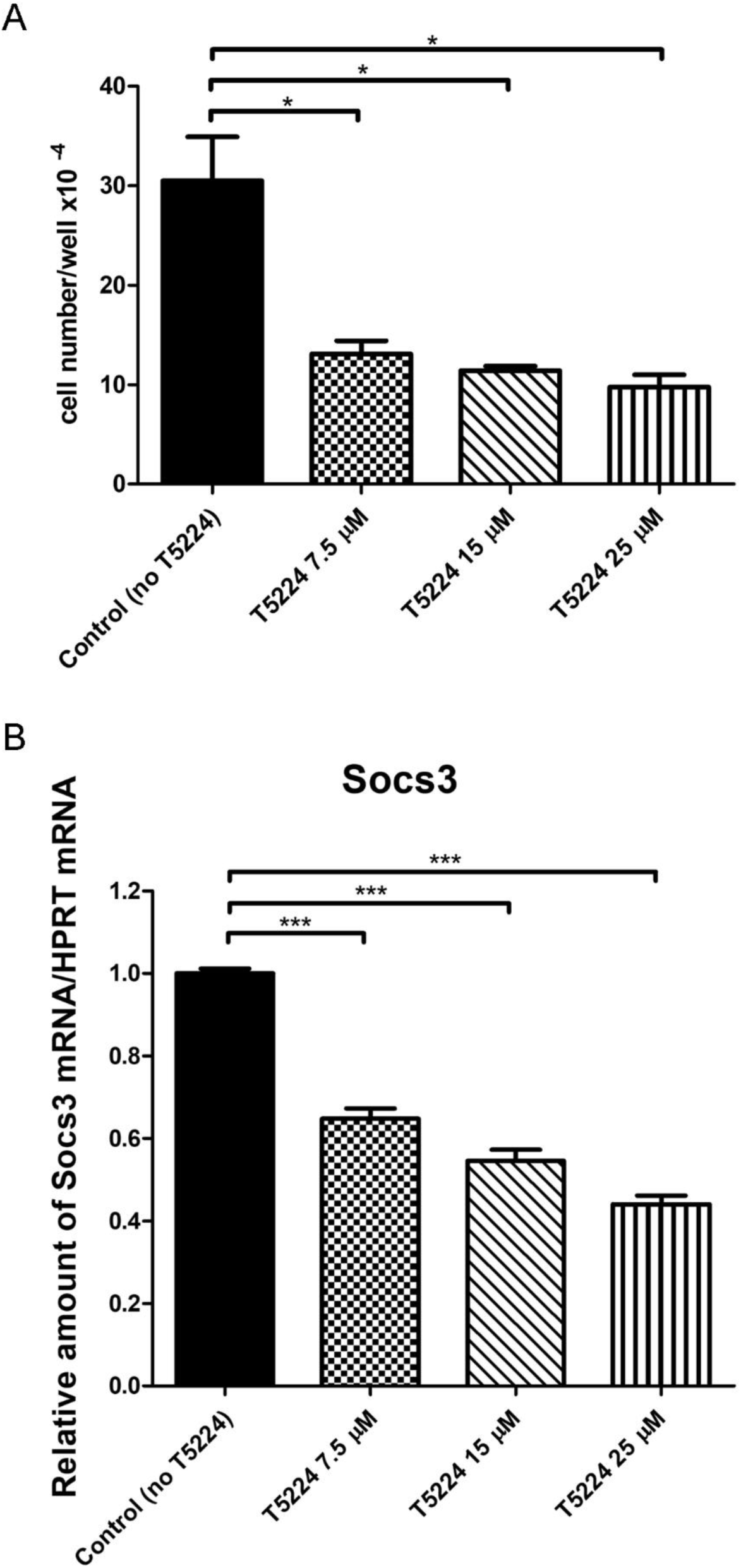
The FOS/JUN (AP1)-specific inhibitor T-5224 impairs NSC growth and Socs3 mRNA expression in a dose-dependent way. **A**, Cell numbers counted 3 days after the administration of T-5224 (dissolved in DMSO) at the indicated concentrations. Control (no T-5224): cells treated with 10 microliters of DMSO only. T-5224: cells treated with T-5224, dissolved in 10 µl DMSO, to obtain the indicated final concentrations of T-5224. Histograms represent the mean ± SEM of results obtained from 3 independent experiments. *p<0.05, paired t-test. **B**, Socs3 mRNA expression relative to HPRT, by real-time qRT-PCR, in cells treated with T-5224, as in A. The values obtained for cells treated with DMSO only (Control (no T-5224)) are set = 1, and the values obtained for cells treated with the indicated concentrations of T-5224 are normalized to the control value. The mean of the absolute values obtained for Control Socs3 mRNA/HPRT mRNA is: 1.051. Histograms represent the mean of results ± SEM obtained with N independent experiments, each analyzed in triplicate by qRT-PCR (***p<0.001, paired t-test): N=3 experiments for T5224 concentrations of 0, 7.5 and 15 µM; N=2 experiments for 25 µM T-5224.

**Figure 5.**
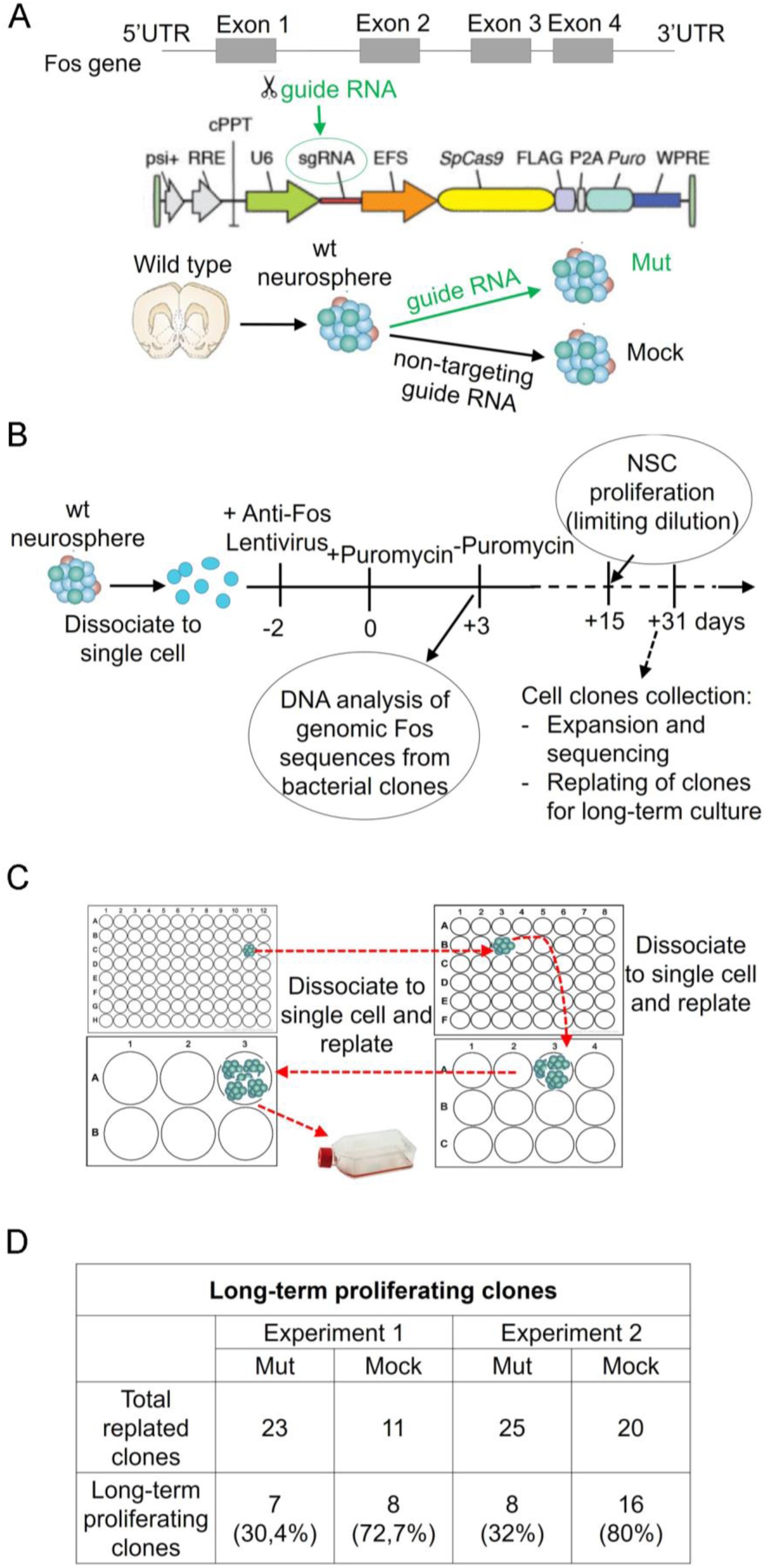
Fos mutagenesis impairs the clonogenic ability of NSC. **A**, Position of the guide RNA-recognized sequence in the Fos gene; lentiviral vector for mutagenesis expressing the guide RNA and Cas9 **B**, Fos mutagenesis experiment and obtainment of transduced, puromycin-resistant NSC as single clones at limiting dilution (primary clones) **C**, Replating of primary single clones for progressive expansion into long-term growing NSC populations (long-term proliferating clones). **D**, Fos mutagenesis in wild type NSC reduces the percentage of long-term proliferating clones. The significance of the difference between number of clones obtained with anti-Fos (Mut) and scramble (Mock) sgRNA was validated by paired one-tailed t-test (p=0.03127).

To verify the efficiency of Fos mutagenesis, at the beginning of the experiment (see above and Fig. 5B) an aliquot of the puromycin-resistant cells was collected, DNA was extracted and amplified by PCR with primers surrounding the site of hybridization of the sgRNA. The amplified DNA was cloned in pGEM®-T Easy (Promega, A1360) by transformation of TOP10 E.coli, and inserts from individual colonies were sequenced. The sequences from CRISPR-Cas9-treated cells were compared to wild-type sequences, by using BLAST NCBI.

Additionally, a number of cell clones obtained at limited dilution were amplified and the sequence of the Fos gene region surrounding the site of the hybridization of the sgRNA was determined.

PCR primers:

Fos Forward: 5’-TCACAGCGCTTCTATAAAGGC-3’ and 5’-CTACTACTCCAACCGCGACT-3’

Fos Reverse: 5’-CTGCGAGTCACACCCCAG-3’ and 5’-CGCCAGTCTCCCTCCAGA-3’

### Immunocytochemistry

Transduced Sox2-deleted, or wild type control NSCs, were dissociated to single cells and seeded on Matrigel™-coated glass coverslips. After 4h cells were fixed (20 minutes) with 4% PFA in phosphate-buffered saline (PBS; pH 7.4) and rinsed 3 times with PBS. Coverslips were then incubated for 90 minutes at 37°C in PBS containing 10% normal goat serum (NGS), 0.2% Triton X-100. Coverslips were incubated with the primary anti-SOX2 antibody (mouse monoclonal IgG2a, 1:100, R&D Systems), overnight at 4°C. Following thorough washing with PBS, cells were incubated for 45 minutes (room temperature) with secondary rhodamine (TRITC)-conjugated goat anti-mouse IgG2a antibodies (1:1000, AlexaFluor Life Technologies). Coverslips were rinsed 3 times in PBS and mounted on glass slides with Fluoromount (Sigma) with DAPI (4′,6-diamidino-2-phenylindole).

#### Pharmacological inhibition of the FOS/JUN AP1 complex

Wild-type NSCs were dissociated to single cells, as described above, and seeded at a density of 1×10^5^ cells/2ml/well in 6-well plates in FM with EGF. 5 mg of T-5224, c-Fos/activator protein (AP-1) inhibitor (MedChemExpress), were dissolved in DMSO according to manufacturer’s instructions to obtain the 5 mM. stock solution. After 4h, cells were treated with T-5224 at concentrations of 7.5 µM, 15 µM and 25 µM. The total DMSO volume added to each culture was adjusted to10 µl in all T-5224-treated samples and in the (DMSO only) control (final DMSO concentration 0.5%). After 3 days, cells were dissociated to single cells, as above, counted, and lysed in TRIzol (Life Technologies) for total RNA extraction.

#### CUT&RUN

Wild-type neurospheres were harvested by pelleting at 180 x g for 4 minutes. CUT&RUN was performed according to [17]. Supernatant was discarded and cells were resuspended in 1 mL room temperature wash buffer (20 mM HEPES pH 7.4, 150 mM NaCl, 2 mM Spermidine, EDTA-free protease inhibitor) by gently pipetting up and down 50 times. Cells were then pelleted twice at 600 x g and resuspended in 1 mL wash buffer after each centrifugation. Cells were counted using an automated cell counter (BioRad), and 100 000 cells per antibody were transferred to a fresh 2 mL tube. Sample volume was adjusted to 1 ml using wash buffer, before 10 µl of ConA bead slurry (prepared according to [17]) per antibody was added to the tube. Cells were incubated with beads on a rocking table at room temperature for 10 minutes, before the tube was placed on a magnetic rack to collect beads. After removing the supernatant, cells were removed from the rack and resuspended in a total volume of 150 µl per antibody of antibody buffer (20 mM HEPES pH 7.4, 150 mM NaCl, 2 mM Spermidine, EDTA-free protease inhibitor, 0.025% Digitonin, 2 mM EDTA) kept on ice. Total volume was split evenly into fresh tubes, one for each antibody. 1.5 µl antibody was added to each sample and tubes were placed on a rocking table in a cold room (4°C) overnight. Cells were collected on a magnetic rack, the supernatant was discarded, and cells were resuspended in 1 ml Dig-wash buffer (20 mM HEPES pH 7.4, 150 mM NaCl, 2 mM Spermidine, EDTA free protease inhibitor, 0.025% Digitonin). The procedure was repeated and cells were collected on the rack again. Supernatant was discarded and replaced with 150 µl of protein A-Micrococcal Nuclease (pA-MNase) fusion protein (700 ng/ml, received as a gift from the lab of Steven Henikoff) in Dig-wash buffer. Cells were incubated on a rocker in a cold room for 1 hour.

Cells were washed two more times by binding beads to rack, removing supernatant, and resuspending in Dig-wash buffer. Supernatant was replaced with 100 µl Dig-wash buffer, gently resuspending beads. Tubes were placed in a heat block kept in wet ice to cool to ca. 0°C. When chilled, 2 µl CaCl_2_ (0.1 mM) was added to each sample. Tubes were mixed by gentle flicking and placed back into block to digest for 30 minutes. Reaction was halted by the addition of 100 µl 2X STOP buffer (340 mM NaCl, 20 mM EDTA, 4 mM EGTA, 0.05% Digitonin, 100 µg/ml RNAse A, 50 µg/ml Glycogen).

The following antibodies were purchased from antibodies-online.com, Aachen, Germany: anti-c-FOS (ABIN2682008), anti-JUN (ABIN3020286), anti-SOX2 (ABIN2855074). A positive control antibody (anti-H3K27me3; (ABIN6923144)) was run with each experiment to validate the technique (data not shown). Anti-HA was purchased from Merck, Darmstadt, Germany (05-902R). Tubes were placed in a 37°C heat block for 30 minutes to release soluble chromatin from cells. Beads were bound to the magnetic rack, and the supernatant was carefully collected into a low DNA binding micro tube. DNA was extracted using Phenol:Chloroform extraction, and libraries were prepared using KAPA HyperPrep Kit (Roche, 07962347001) using KAPA DUI adapters (Roche, 8278555702). Libraries were pooled and sequenced on an Illumina NextSeq 500 sequencer using a NextSeq 500/550 High Output Kit v2.5 (75 Cycles), 36bp PE, generating FASTQ files.

Reads were aligned to the mouse genome (mm9) using bowtie [18] with settings -X 700 -m1-v 1. Reads were trimmed using BBDuk (*BBMap – Bushnell B. – sourceforge*.*net/projects/bbmap/*), removing overrepresented repeat sequences (i.e., [TA] _18_) and adapter sequence. Reads were filtered for size, keeping only reads with a fragment size at or below 150 base pairs. Bedgraph files were generated using bedtools genomecov [19], and peaks were called using SEACR [20], in relaxed mode, normalizing to control.

## Results

### Reactivation of Fos expression in Sox2-deleted neural stem cells restores long-term self-renewal

We identified, by RNAseq, about 900 genes downregulated in NSC grown in vitro from neonatal mouse brain, after Cre-mediated Sox2 deletion during embryogenesis [14]. To identify the network of genes mediating Sox2 function in NSC self-renewal, we reasoned that genes downregulated following Sox2 deletion likely include those required for self-renewal; indeed, we previously showed that the most downregulated gene, Socs3, encoding a signaling protein acting at the cell membrane, rescued long-term self-renewal when expressed via a lentivirus in Sox2-deleted cells [14]. We therefore overexpressed, in Sox2-mutant cells, three among the most downregulated genes (Fig. 1A), Fos, Jun, and Egr2, encoding transcription factors which might possibly be involved in the regulation of Socs3 (or other critical factors).

To test for the functional role of the downregulated genes, we individually cloned their cDNAs into a lentiviral expression vector, and transduced them into Sox2-mutant cells, cultured from whole forebrains of P0 mice from which Sox2 had been deleted in vivo, at embryonic day (E) 11.5, via a nestin-Cre transgene [5,14](Fig. 1B). We first tested a combination of vectors expressing Fos and Jun, since FOS and JUN are known to act together, forming the AP1 complex, in transcriptional regulation [21]. The vector contains a delta-NGF receptor (dNGFR) marker gene that allows to identify transduced cells by anti-dNGFR immunofluorescence (Fig. 1B). In preliminary experiments, we transduced NSC using single vectors, including each of the genes to be tested, to evaluate the proportion of the cells that were transduced by each type of vector: the transduction efficiency was 23,3% for Fos; for Jun and for Egr2, it was 18,5% and 17,1%, respectively.

We then transduced mutant cells at early stages of their culture, using combinations of different vectors at the same multiplicity of infection (MOI) as that used for single-vector transduction (Fig. 2). After initial passages, non-transduced mutant cells progressively slowed down their proliferation and stopped growing, the culture eventually becoming exhausted, as expected [5,14]; by contrast, cells transduced with Fos and Jun continued to grow exponentially, with a kinetics matching that of control wild-type cells (Fig. 2A; Suppl. Fig.1). Taking advantage of the delta-NGF receptor (dNGFR) marker, that is coexpressed with the cDNAs from the lentiviral vectors, we followed the percentage of dNGFR-positive (transduced) cells through cell passaging. While, at the beginning (passage 2 after transduction), dNGFR-positive cells represented about 50% of total cells, their percentage progressively rose to 100% (passage 15) through passaging (Fig. 2C; Suppl. Fig. 2), indicating a selective advantage of transduced cells.

We also verified that cells that had recovered self-renewal were all SOX2-negative, by immunofluorescence (IF) for SOX2 (Suppl. Fig. 3A); this ruled out the possibility that a positive selection of rare, SOX2-positive cells persisting in Sox2-deleted brains could contribute to the recovery of mutant cells.

We also tested a second combination of vectors, expressing Fos and Egr2 (Fig. 2B). In parallel with this vector combination, we also transduced mutant cells with the Fos-expressing vector only (Fig. 2B). The viral upregulation of Fos and Egr2 in mutant cells prevented cell exhaustion, and restored long-term self-renewal (Fig. 2B), similarly to what had been found with Fos+Jun (Fig. 2A). A similar result was obtained also with the Fos vector alone; self-renewal was restored, and the growth kinetics of the cells was similar to that of wild type cells (Fig. 2B; Suppl. Fig. 1). We then measured the percentage of transduced cells by FACS analysis with an anti-dNGFR antibody, as previously done for the Fos+Jun-transduced cells; this was about 20% at the beginning (passage 2), and progressively rose to about 100% through passaging (for both Fos+Egr2 and Fos only), again pointing to a selective advantage of the transduced cells (Fig. 2C; Suppl. Fig. 2). As a control experiment, we also transduced wild type cells with the Fos-expressing vector; this did not lead to any change in the growth kinetics of the cells, that remained the same as that of non-transduced cells (Suppl. Fig. 3B). As a further control, we also verified that mutant cells, transduced with an “empty” dNGFR lentiviral vector, did not change their growth kinetics in comparison to that of non-transduced cells (Suppl. Fig. 3B).

Overall, these results point to Fos as the major player in the rescue of NSC self-renewal in transduced cells.

Do any of the other co-transduced cDNAs play any additional role in the effect shown? To this end, we plated the transduced cells (Fos + Jun transduction) at limiting dilution, in 96 well plates, to obtain clones. These were then individually expanded and tested for the presence of the two lentiviral inserts (Fos, Jun). Out of 14 clones tested by PCR analysis, only two contained Jun, but all contained the Fos-expressing vector (Fig. 3A). This indicated that Fos is required for the observed recovery of self-renewal, as suggested. Importantly, no clones exclusively transduced with Jun (but not with Fos) were detected, implying that Jun, by itself, is not sufficient to rescue Sox2-deleted stem cells. The proportion of doubly transduced clones (Fos+, Jun+) is roughly that expected on the basis of the probability of chance transduction of a Fos-transduced cell also with a Jun vector, based on the previous assessment of the transduction efficiency of individual vectors (see above).

Similarly, in the Fos+Egr2s-transduction, out of 14 clones expanded and analyzed, 8 contained the Egr2 vector, but all contained the Fos vector; none contained the Egr2 vector alone (Fig. 3B). Again, this indicates that Egr2 alone cannot rescue NSC long-term proliferation.

#### Pharmacological inhibition of the FOS/JUN AP1 complex antagonizes NSC growth and reduces Socs3 mRNA levels

FOS is transcriptionally active as an heterodimer (AP1 complex) with JUN, or another JUN-like factor [22]. Both Fos and Jun mRNAs are strongly down-regulated in Sox2-deleted NSC (Fig. 1). T-5224 is a drug capable of binding to the interface of FOS and JUN in the AP1 complex, thus selectively inhibiting AP1 binding to the cognate DNA sequences [23,24]. We reasoned that treatment of wild-type NSC with T-5224 might somehow mimic the observed FOS and JUN decrease in Sox2-deleted NSC, giving information on the role of FOS (and JUN) in NSC. We thus treated wild type NSC with T-5224; the treatment reduced cell proliferation, after 3 days, by 50-60%, at concentrations of 7.5-25 µM (Fig. 4A), comparable to concentrations found to inhibit luciferase-reporter activity guided by AP1, but not by other transcription factors [23]; higher concentrations (40-80 µM) were previously shown to inhibit T lymphocyte proliferation in vitro [25]. Further, we detected a clear reduction of Socs3 mRNA expression, with significant inhibition (30-55%) between 7.5 and 25 µM T-5224 (Fig. 4B). These results are consistent with the hypothesis that the reduction of Fos and Jun observed in Sox2-deleted NSC affects cell proliferation and expression of the putative Sox2 target Socs3.

#### Fos mutation reduces NSC long-term self-renewal

The decreased NSC proliferation upon FOS/JUN inhibition raised the question whether Fos activity plays a role in the maintenance of long-term NSC renewal, in the presence of wild type Sox2. We thus mutated the endogenous Fos gene, using the CRISPR/Cas9 system. A short guide RNAs (sgRNA) was identified, which efficiently mutagenized Fos (sgRNA-fos) (Fig. 5A; Methods). Wild-type NSC were transduced with a Cas9+sgRNA-expressing lentivirus, carrying a puromycin-resistance gene, or a control virus expressing “scramble” sgRNA that is not expected to mutate Fos; transduced cells were then selected by puromycin for 3 days (Fig. 5B). After puromycin withdrawal, the cells were grown for 15 days to allow recovery from the stress of transduction and selection; at this initial stage, control and mutated cells proliferated in bulk culture with comparable kinetics (not shown). In parallel, at the time of puromycin withdrawal, an aliquot of the puromycin-resistant cells was collected, DNA was extracted and amplified by PCR with primers surrounding the site of hybridization of the sgRNA, and the PCR product cloned in a plasmid, and sequenced (Fig. 5B). Further sequences were obtained using samples taken at day 8 and day 20 after puromycin withdrawal. Of 36 different PCR products analyzed by sequencing, one was wild type, and the remainder were mutated by insertion or deletion (indel) of nucleotides; 10 of the indels still maintained the appropriate reading frame (Supplementary Table 5). We also sequenced cell clones obtained from propagation of single cells (see below): of 9 clones sequenced, 1 was wild type, 2 were heterozygous mutants, and 6 were homozygous mutant clones: note that several clones died in the early stages of expansion, and could not be sequenced, suggesting that some selection against mutated cells had occurred. In contrast, in scramble-treated cells no mutated Fos gene was found (Suppl. Table 1). Finally, we checked sequences bearing partial complementarity to the sgRNA target region in Fos, to ascertain the potential presence of “off-target” mutations, which might affect genes potentially controlling NSC self-renewal. We used a similar strategy to [26,27]; we identified, by an informatic genome-wide survey using GT-scan [28], 6 regions (called A to F) carrying an appropriately located PAM sequence, and bearing similarity to the sgRNA used. All these sequences had 3 mismatches with the sgRNA (Suppl. Table 3); we did not find any sequence with 1 or 2 mismatches. By cloning PCR-amplified regions A to F into plasmids, and individually sequencing them, we showed that only region D showed one mutation (in 1 out of 21 different samples sequenced) (Suppl. Table 4). Overall, these data show that CRISPR-Cas9 mutagenesis of the Fos gene was efficient and free of frequent off-target mutations in partially complementary regions.

We then tested the ability of NSC to self-renew generating stem cells capable of giving rise to clonal progeny [29,30](Fig. 5C). In this type of experiment, cells were plated at clonal dilution in individual wells, and the appearance of single neurospheres in a well was evaluated at the microscope. When the growth of the neurosphere had generated a relatively large number of cells (a “primary clone”), this was picked up, disaggregated and replated (“replated clones”, Fig. 5C,D) in a single well, for further expansion. Primary clones could be obtained from both scramble-treated controls (mock) and sgRNA-Fos treated cells. We then asked whether each of these clones represents the bona fide progeny of a long-term self-renewing stem cell, by testing for its ability to expand through serial passaging, to give long-term growth (Fig. 5C,D Suppl. Table 2, “long-term proliferating clones”). We performed two experiments (Fig. 5D, Suppl. Table 2): in both experiments, the number of primary clones (from sgRNA-fos treated cells), capable to reform secondary clones demonstrating long-term growth, was greatly, and significantly, diminished in comparison to controls from mock-treated cells (Fig. 5D, Suppl. Table 2, line 2). We note that the potential off-target region D showed a point mutation in 1 out of 21 individual sequences from Fos-mutated cells, lying in the 5’ UTR of the Max gene (Suppl. Table 4). While it is possible that this mutation might affect translation of the mutated Max mRNA, there is no immediately obvious mechanism for such an effect . More importantly, our study of long-term self-renewal of Fos mutated cells analyzed about 50 individual clones; it is highly unlikely that this, or other very low frequency mutations (if any), significantly contribute to the strong decrease of the number of long-term proliferating Fos-mutant clones (versus wild-type clones) observed in the two experiments performed.

Overall, these data (Fig. 4 and Fig. 5). indicate that FOS-deficient NSC progressively deteriorate during culture, failing to self-renew, a phenomenon reminiscent of the previous observations obtained with Sox2-deleted NSC [14].

#### Sox2 upregulation in Sox2-mutant cells restores expression of endogenous Fos and Socs3

These observations made us ask if re-expression of Sox2 into Sox2-deleted cells [5](Suppl. Fig. 4), would also re-increase levels of Fos and Socs3 expression. As expected, Sox2 re-expression in Sox2-deleted NSC fully recovered normal long-term self-renewal (Suppl. Fig. 4); by contrast, Empty Vector transduction is unable to rescue long-term self-renewal of Sox2-deleted NSC ([31] and data not shown). A selective advantage of Sox2-transduced cells was again observed by FACS analysis through passaging, until almost 100% of the cells were represented by transduced cells (Suppl. Fig. 4).

We thus analyzed by qRT-PCR the expression of Fos, and of Socs3, in mutant cells (three different mutants) transduced with Sox2, which re-acquire the ability to self-renew long term. Transduction of Sox2 initially (until passages 5-6) had little or no effect on the expression of Fos and Socs3 relative to control un-transduced cells (Fig. 6A,B), and to empty vector transduced cells (not shown); however, at later stages in culture, after passage 6, there was a substantial increase of both Fos and Socs3 mRNAs (relative to the levels observed in the first 5 passages) that lasted at least until late passages (passages 20-22, when the experiment was interrupted)(Fig. 6A,B). In some experiments, we also checked the expression of Socs3 in Sox2-deleted NSC transduced with Fos (alone or together with Jun or Egr2); again, a late increase of Socs3 expression, versus Empty Vector-transduced (not shown) or un-transduced NSC, was observed (Fig. 5C).

**Figure 6.**
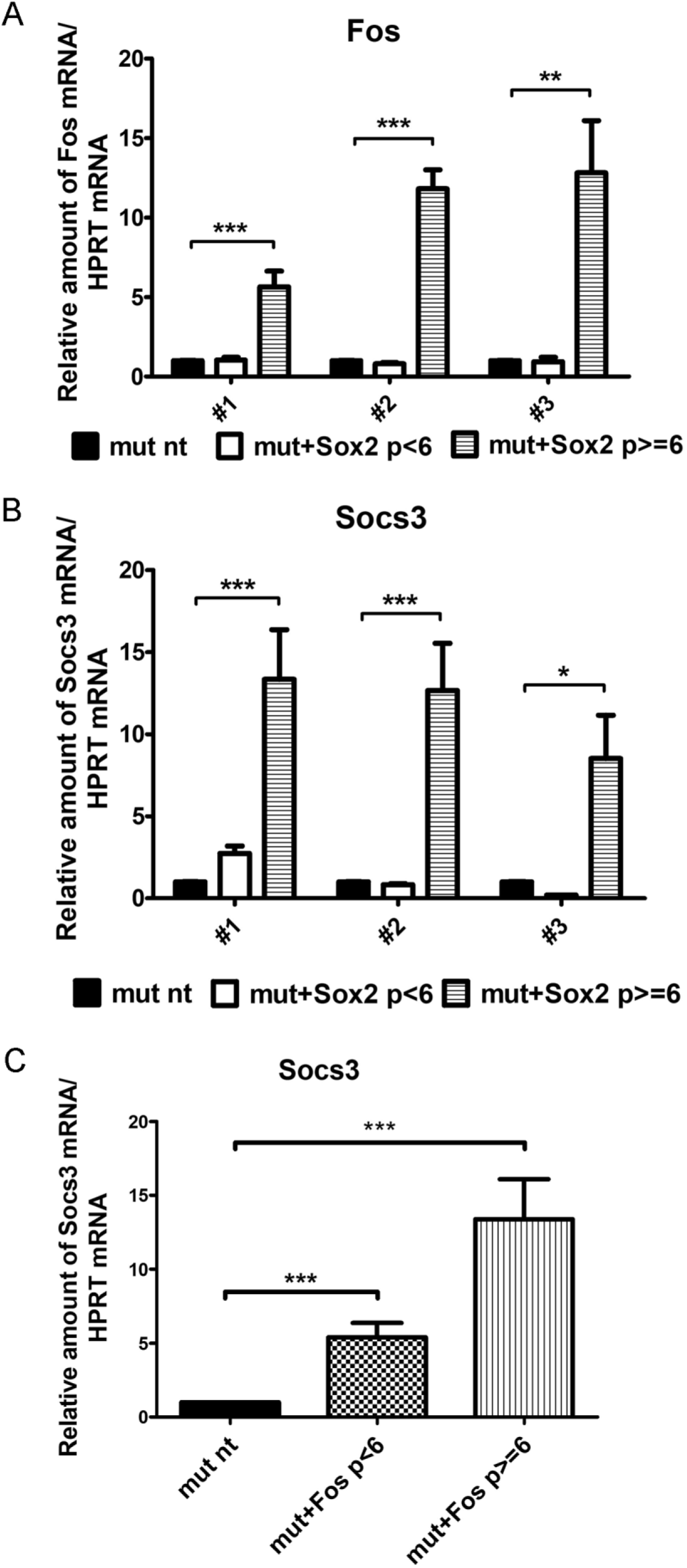
Sox2-rescued Sox2-deleted NSC recover expression of endogenous Fos and Socs3. **A**,**B**, Expression of Fos (A) and Socs3 (B) in Sox2-deleted NSC(three different mutants, #1, #2, #3) transduced with Sox2-expressing lentivirus. P<6 and p>6 are the average expression values for passages 2-5 and 6-22, respectively. **C**, Expression of Socs3 in Sox2-deleted cells transduced with Fos. mut: Sox2-deleted. nt: non transduced. In A,B and C, values reported for non-transduced Sox2-deleted NSC (mut nt) were obtained at the initial passages 1-3, when cells are still actively growing. In A,B,C, expression values are calculated as ratios of Fos or Socs3 mRNA/HPRT mRNA, as determined by qRT-PCR. The values obtained for untransduced mutant NSC (mut nt) are set = 1, and the values obtained for Sox2- or Fos-transduced mutants (mut + Sox2; mut + Fos) are normalized to the corresponding mut nt. The mean of the absolute values obtained for mut nt Fos mRNA/HPRT mRNA are: 25.30 (#1), 16.38 (#2) and 14.63 (#3). The mean of the absolute values obtained for mut nt Socs3 mRNA/HPRT mRNA are: 2.30 (#1), 1.43 (#2) and 1.99 (#3). Histograms in A,B,C represent the mean of results obtained with the following numbers of qRT-PCRs at different passages: mut 1 and 2: 20 qRT-PCRs performed in duplicate for mut nt and mut+Sox2 p>6, and 13 qRT-PCRs performed in duplicate for mut nt and mut+Sox2 p<6; mut 3: 10 qRT-PCRs for nt and mut+Sox2 p>6; 3 qRT-PCRs for mut nt and mut+Sox2 p<6. ***p <0.001; *p<0.05, paired t-test.

#### Both SOX2 and FOS are direct regulators of Socs3 expression

Previously, we showed that Sox2 binds to the Socs3 gene promoter in NSC chromatin, as well as to a distant enhancer connected to the promoter by a long-range interaction; experiments in zebrafish proved the in vivo functional activity of the enhancer, in the forebrain [14]. Fos binding to the Socs3 promoter and its functional activation of the gene were previously demonstrated by gel-shift experiments in non-neural cells [32,33]. It remained to be determined whether the association between FOS and the Socs3 promoter could also occur in NSC chromatin in vivo. To test this, we used the sensitive CUT&RUN procedure[17,20], which allows detection of the genomic regions occupied by chromatin-binding proteins in living cells. that detects chromatin-bound proteins by cleavage of the DNA adjacent to the bound protein by pA-MNase bound to a specific antibody. Fig. 7 shows that both FOS and JUN, the two components of AP1, are bound to a region overlapping the Socs3 promoter, and corresponding to the SOX2 peak. Consistently, a modified AP1 consensus site was detected in that region by previous investigators [32]. These results indicate that Socs3 is not only a target of SOX2 but also of the downstream effectors FOS and JUN, suggesting the existence of a complex SOX2-driven gene regulatory network required to maintain stemness properties in NSC.

**Figure 7.**
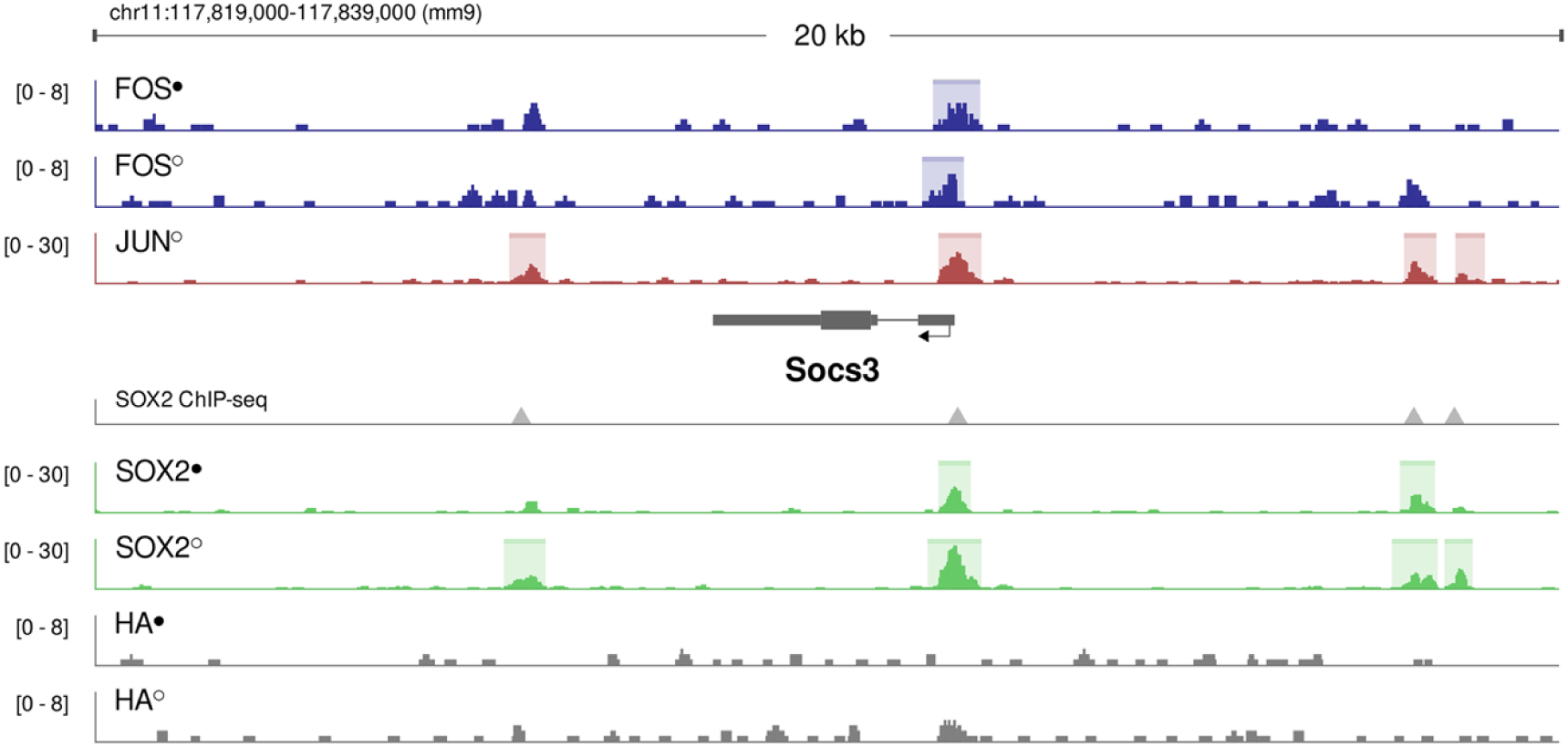
CUT&RUN detects FOS, JUN and SOX2 binding onto the Socs3 promoter in NSC chromatin. CUT&RUN targeting different transcription factors in neurosphere cells. CUT&RUN was performed on 100,000 neurosphere cells per antibody. Tracks show a duplicate experiment (replicates marked with • or ○), one of which also shows an anti-JUN antibody. Peaks called by SEACR [20] are shown as colored boxes. Location of peaks called using ChIP-seq targeting SOX2 in neurospheres [14] are indicated as grey triangles.

## Discussion

### Fos acts as a mediator of Sox2 function in NSC self-renewal in vitro

The transcription factor Sox2 regulates a downstream gene expression program necessary to maintain the ability of NSC to self-renew [5]. A critical Sox2-target gene is Socs3: indeed, Socs3 overexpression is able to rescue the self-renewal defect of Sox2-mutant NSC [14]. Do any additional genes, upstream to Socs3, or acting in parallel with it, contribute to mediate Sox2 effects on self-renewal? In the present work, we tested some candidate genes among those which are more strongly downregulated in Sox2-mutant cells. We found that Fos alone is able to fully rescue Sox2-mutant cells (Fig. 2). Clonal analysis (Fig. 3) shows that all clones obtained after rescue of long-term proliferation by Fos transduction contain the transduced Fos gene; as the initially transduced cells represent a minority in the population, this finding indicates that the rescue is due to positive selection of Fos-transduced cells, and not by (rarely occurring) chance events. The analysis of the rescued clones, obtained by Fos+Jun or Fos+Egr2 transduction, shows that neither Jun, nor Egr2 play critical roles in the rescue. In fact, we do not find any rescued clones that have been transduced exclusively with one of these genes, and, when these transduced genes are present in clones, they are always accompanied by Fos. Further, the proportion of Fos-transduced clones, that are also transduced by Jun (Fos + Jun), is not higher than that expected on the basis of chance cotransduction of a Fos-transduced cells also by a Jun vector, indicating that Jun does not significantly cooperate with Fos in the rescue (Fig. 3). However, the proportion of Fos/Egr2 cotransduced rescued clones (8 out of 14 total rescued clones) is higher than expected based on the assumption that Fos transduced cells should also be transduced with Egr2, by chance, with a probability of 0.17, the frequency of Egr2 transduction; the result obtained might suggest that Fos-transduced cells, also transduced with Egr2, have some proliferative advantage over Fos-only-transduced NSC, indicating that Egr2 could cooperate with Fos.

We wished to confirm the evidence that Fos is important for long-term self-renewal of NSC by performing the reciprocal experiment, i.e. by inhibiting the activity of FOS in cells carrying intact Sox2. The significant inhibition of NSC proliferation observed upon treatment with the T-5224 inhibitor of AP1 activity (Fig. 4) is in agreement with an important role of FOS for the proliferation of NSC. In addition, the progressive decrease of clonogenicity that occurs in Sox2-positive NSC mutated in Fos by CRISPR/Cas9 (Fig. 5; Suppl. Tables 1, 2) extends these findings to the concept that Fos is a relevant component of the genetic program controlling in vitro long-term self-renewal of NSC.

Fos, a component, with Jun or related factors, of the AP-1 protein complex, is well known to control cell life and death by regulating the expression and function of cell cycle regulators such as Cyclin D1, p53, p21cip1/waf1, p19ARF and p16 [22]. In NSC grown in vitro, factors such as FGF2 and bEGF control cell proliferation through a cascade involving ERK signaling, Fos, and cyclins [34]. The strong decrease of Fos in Sox2-deleted cells (Fig. 1A), although less profound than the Socs3 decrease, is in line with the known role of Fos in cell proliferation control. Thus, five lines of evidence support the conclusion that Fos is an important mediator of Sox2 function in the maintenance of NSC self-renewal: a) the high abundance of Fos mRNA in wild type NSC, and the strong decrease of its level in Sox2-deleted cells; b) the rescue of self-renewal of Sox2-deleted cells by Fos overexpression (which, by itself, has no effect in wild type NSC); c) the inhibition of NSC proliferation by T-5224, a transcriptional inhibitor of the FOS/JUN complex; d) the requirement for Fos for efficient long-term NSC self-renewal; e) the known proliferation-related activity of Fos in several different cell types .

In conclusion, Fos appears to be part, together with Socs3, of the Sox2-dependent gene expression program regulating NSC maintenance. We note that there may be, in theory, several additional Sox2-regulated genes critically involved in NSC self-renewal, and some of them might act in a complementary way; the overexpression strategy used here may be exploited to identify them. On the other hand, we do not expect that the single removal of the Fos (or Socs3) gene would necessarily lead to complete NSC loss, in vitro and in vivo. Indeed, in our Fos mutagenesis experiments, single cells-derived primary clones do arise and some of them are able to establish secondary cultures, some of which can be further expanded into cell populations (Fig. 5). This is in agreement with a substantial conservation of overall neural cell proliferation during development reported in Fos-knock-out mice [35]. Whether Fos knock-out in vivo would cause important defects in NSC survival/proliferation, pre-and post-natally, as observed in Sox2 mutants [5,36], remains to be investigated.

### Defining a functional gene regulatory network acting downstream to Sox2 in NSC self-renewal

Sox2 re-expression in Sox2-deleted cells is able to efficiently rescue long-term NSC proliferation, similarly to the effects observed upon Socs3 [14] or Fos transduction of Sox2-deleted cells. Is there a direct relationship between Sox2 activity and Fos and/or Socs3 expression? Genomic evidence from [14], and from the present paper (Fig. 7), suggests that this may be the case. In favor of this hypothesis is the observation that Fos is connected by long-range chromatin interactions to two different functionally validated enhancers [14,37], shown by SOX2 ChIPseq to be bound by SOX2 (see Table S7 in [14]), and that Socs3 is bound by SOX2 on its promoter (Fig. 7) as well as on a connected functional enhancer [14]. Additionally, also Jun and Egr2 gene regions show SOX2 binding (Suppl. Fig. 5). Sox2 loss disrupts a large proportion of promoter-enhancer interactions in NSC, including those in the Fos, Socs3, Jun and Egr2 regions (Suppl. Fig. 5), providing a plausible explanation for the decreased expression of all these genes detected by Bertolini et al [14], see also Fig. 1.

Finally, additional studies in non-neural cells showed that both the AP1 (FOS+JUN) complex and Egr1/2 interact with the Socs3 promoter to activate it in transfection experiments (Fig. 8) [32,33,38]. In addition, impacting Fos activity via AP1 inhibition by T-5224 treatment (Fig. 4) leads to decreased Socs3 expression (Fig. 4). In agreement with these data, we now show that both FOS and JUN bind, together with SOX2, to the Socs3 promoter region (Fig. 7).

**Figure 8.**
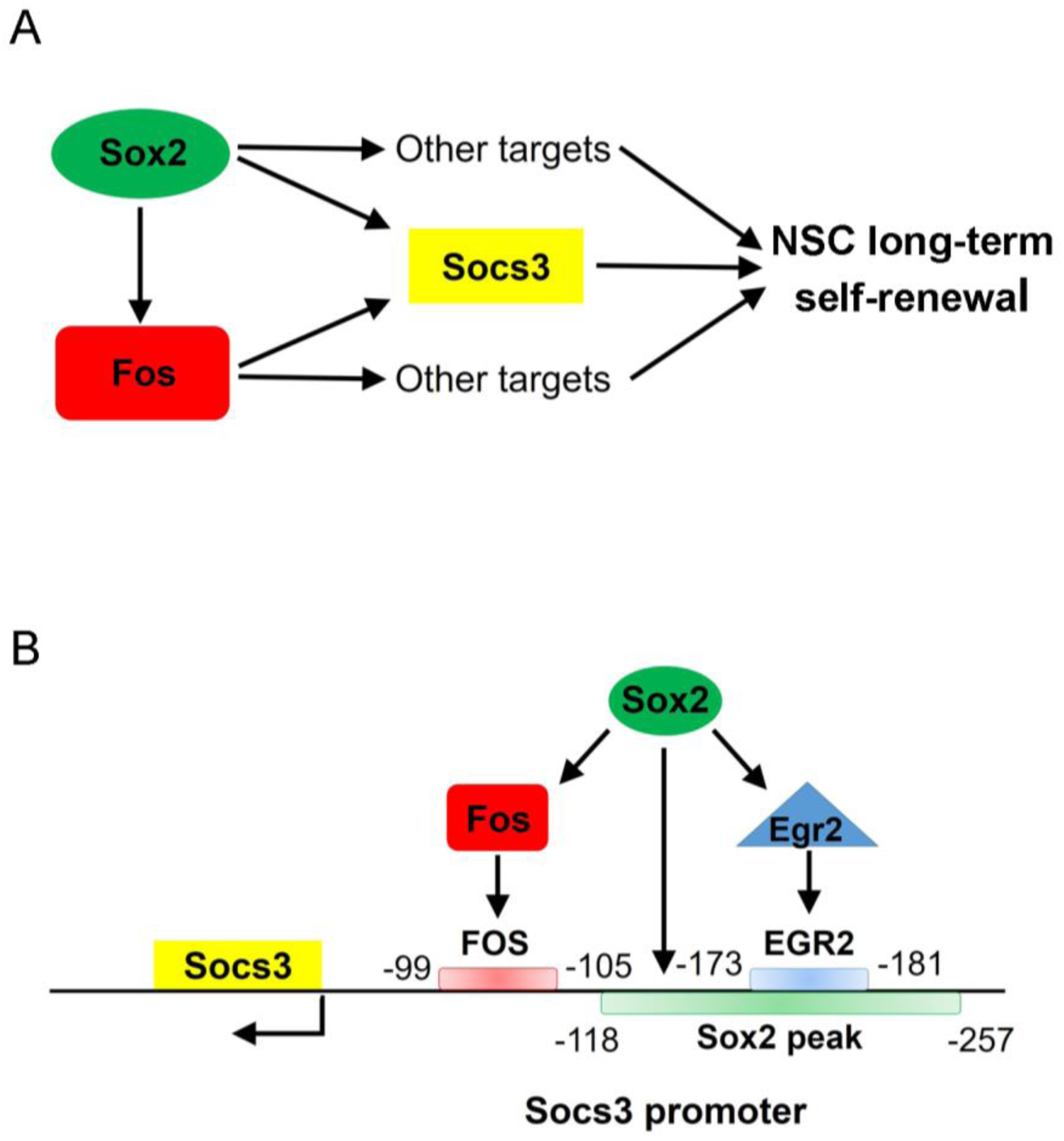
A model for a Sox2-controlled gene regulatory network involving Fos, Egr2 and Socs3. **A**, Regulatory relations between Sox2, Fos, and Socs3, in NSC long-term self-renewal control **B**, Sox2, Fos and Egr2 functional interactions on the Socs3 promoter. For Fos and Egr2 binding data, see [32,38]. For Sox2 binding, see [14].

As to Egr2, preliminary experiments (not shown) suggest that Egr2 transduction in Sox2-deleted cells transiently increases Socs3 levels during the initial cell culture passages. On the other hand, it should be remarked that Sox2 transduction affects Fos and Socs3 gene expression only at relatively late stages (after passage 6, Fig. 6). We speculate that several genes, downstream to Sox2, and related signaling pathways, have to be activated in concert in order to re-establish proper levels of expression of Fos and Socs3. Fos and Socs3 expression are also known to be critically dependent on signals emanating from many different growth factors [22,24,32,33,39]. Moreover, re-establishing the correct 3D structure of chromatin following its disruption due to Sox2 deletion, may require a relatively long time. It may be noted, in this context, that reprogramming of somatic cells into iPS cells by OSKN genes, of which Sox2 is part, typically requires extended lengths of time.

### Conclusions and Summary

Overall, these data indicate that a Sox2-dependent Fos/Jun/(Egr)-Socs3 network participates in the regulation of in vitro NSC self-renewal (Fig. 7,8). Within this network, Fos and its putative target Socs3 appear to be essential for reestablishing NSC renewal in Sox2-deleted NSC. While our transduction results do not document a critical role for the Jun and Egr genes in NSC maintenance, their proposed involvement in Socs3 regulation (Fig. 8), their dependence on Sox2 for expression, and their binding by Sox2 (Suppl. Fig. 5) point to some role of these genes in Sox2-dependent proliferation. Indeed, it remains possible that the overall residual levels of Jun and Egr1/2 in Sox2-deleted cells are sufficient (and required) to sustain their regulatory functions in NSC, once Fos is provided to the cells by transduction, as in Fig. 2. Under these conditions, further adding Jun and Egr expression, as in Fig. 2, is not expected to reveal their necessity. The present data open a new window on factors necessary for NSC maintenance.

## Acknowledgements

We thank A. Ronchi for providing the pHR SIN BX IR/EMW vector. This research was supported by ERANET – NEURON ImprovVision and Associazione Italiana per la Ricerca sul Cancro (AIRC) grant IG 2014 – 16016 to S.K.N. M.P. is the recipient of a Dipartimenti di Eccellenza fellowship. C.B. is the recipient of a DIMET (Doctorate in Molecular and Translational Medicine) PhD fellowship. C.C. is a fellow of the Wallenberg Center for Molecular Medicine and receives generous financial support from the Knut and Alice Wallenberg Foundation, and by Cancerfonden (CAN2018/542). The authors thank the National Supercomputer Centre (NSC-SNIC) at Linköping University for storage and computing resources, and Steven Henikoff and Terri D. Bryson for sharing CUT&RUN reagents.

## Data availability

The data that support the findings of this study are available from the corresponding author upon reasonable request.

