## Supplementary Figures and Table for "Sox2 controls neural stem cell self-renewal through a Fos-centered gene regulatory network"

**Supplementary Figures and Tables**

**Supplementary Figure 1**

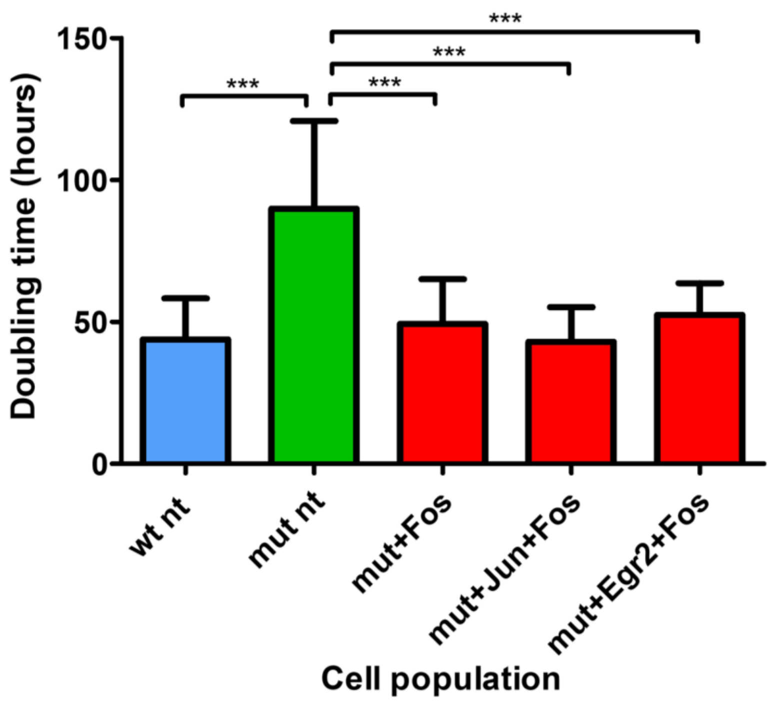


**Supplementary Figure 1**

**Doubling times of wild type, Sox2-deleted, and Sox2-deleted, Fos-transduced NSC in long-term culture**

Population doubling time (hours) of Sox2-wt NSC, Sox2-deleted NSC and Sox2-deleted NSC transduced with Fos alone, Fos+Jun and Fos+Egr2. Sox2-deleted NSC transduced with Fos alone, Fos+Jun and Fos+Egr2 (red) duplicate faster than non-transduced Sox2-deleted NSC (mut nt, green) and their doubling time is similar to that of Sox2-wt NSC (blue). Histograms represent the mean of results ± SD obtained with passages in culture (up to passages 16-20) for Sox2-wt NSC and for Sox2-deleted NSC transduced with the indicated vector, and with 4-7 passages in culture for Sox2-deleted NSC, i. e. up to the last passage before the “plateau” in the growth curve is reached (***p<0.001, unpaired t-test): N = 4 wt nt; N = 4 mut nt; N = 2 mut+Fos; N = 1 mut+Fos+Jun and N = 1 mut+Fos+Egr2.

**Supplementary Fig. 2**

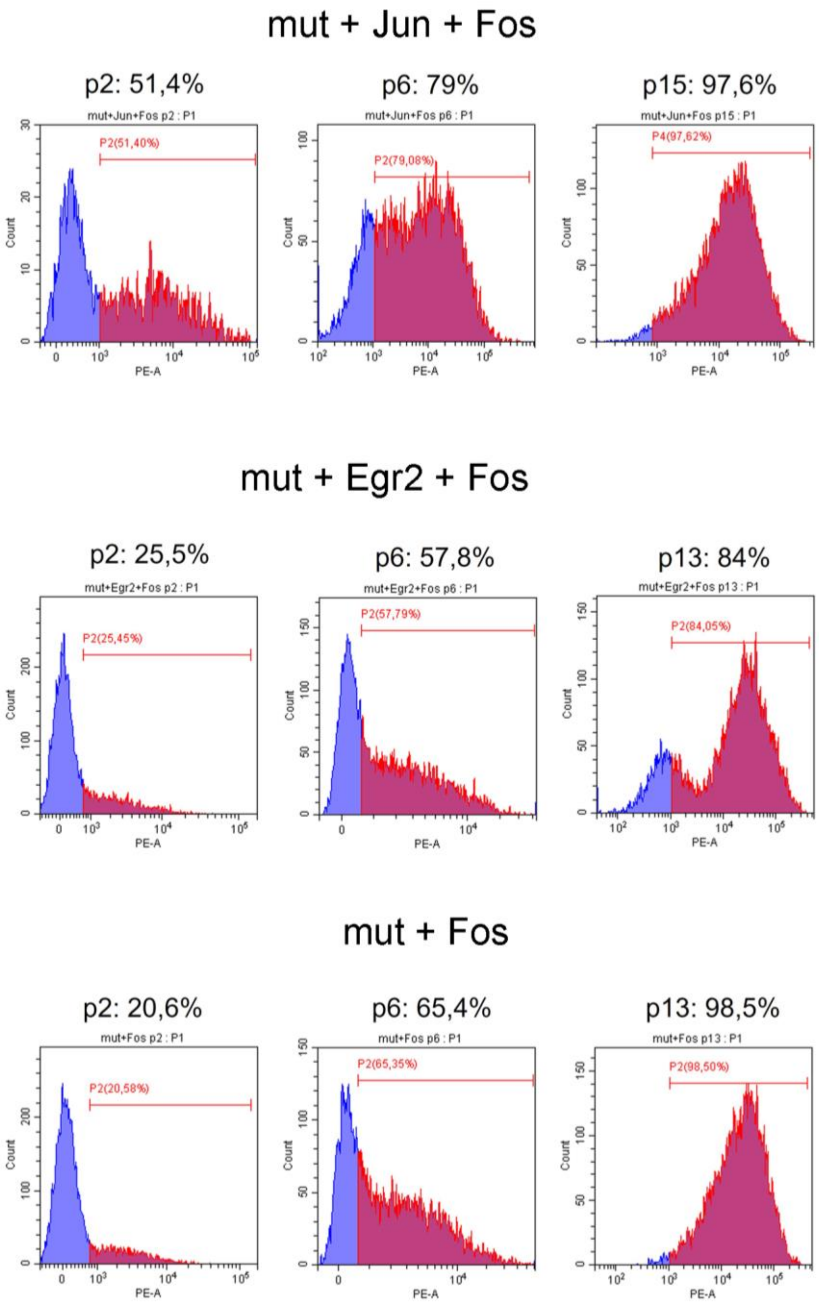


**Supplementary Fig. 2**

**FACS analysis of dNGFR-positive Sox2-deleted cells transduced with Jun and Fos, Egr2 and Fos, or Fos only**

dNGFR-positive cells are indicated by the red color, and the corresponding percentage is indicated. Their proportion increases progressively at different passages.

**Supplementary Fig. 3**

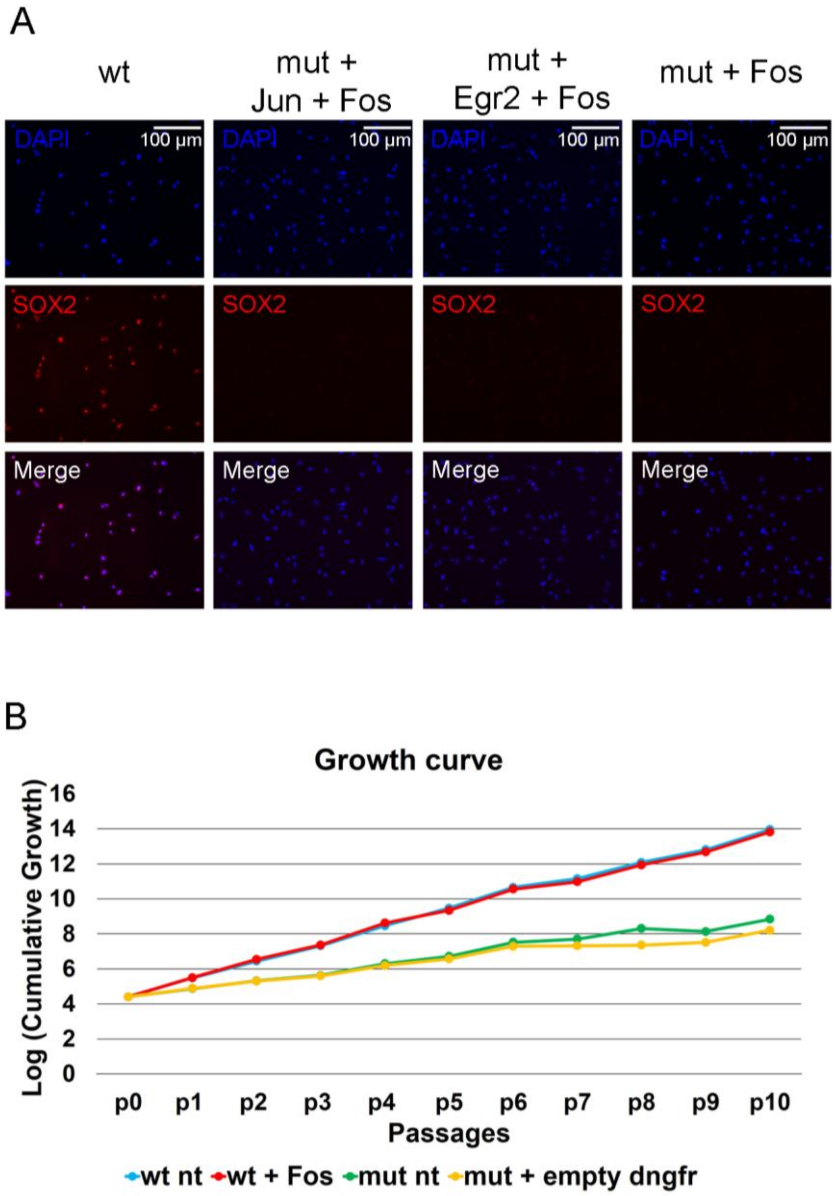


**Supplementary Fig. 3**

**A, Absence of residual SOX2-positive cells in long-term cultures of Sox2-deleted NSC transduced with Fos+Jun, Fos+Egr2, or Fos only**

**B, Growth of wild type NSC transduced with Fos or not transduced (nt). Growth of Sox2-deleted (mut) cells transduced with empty vector (empty dngfr) or not transduced (nt).** One representative experiment out of two is shown for wt nt, wt + Fos, mut nt and mut + empty dngfr. The results of the replica experiments are essentially superimposable.

**Supplementary Fig. 4**

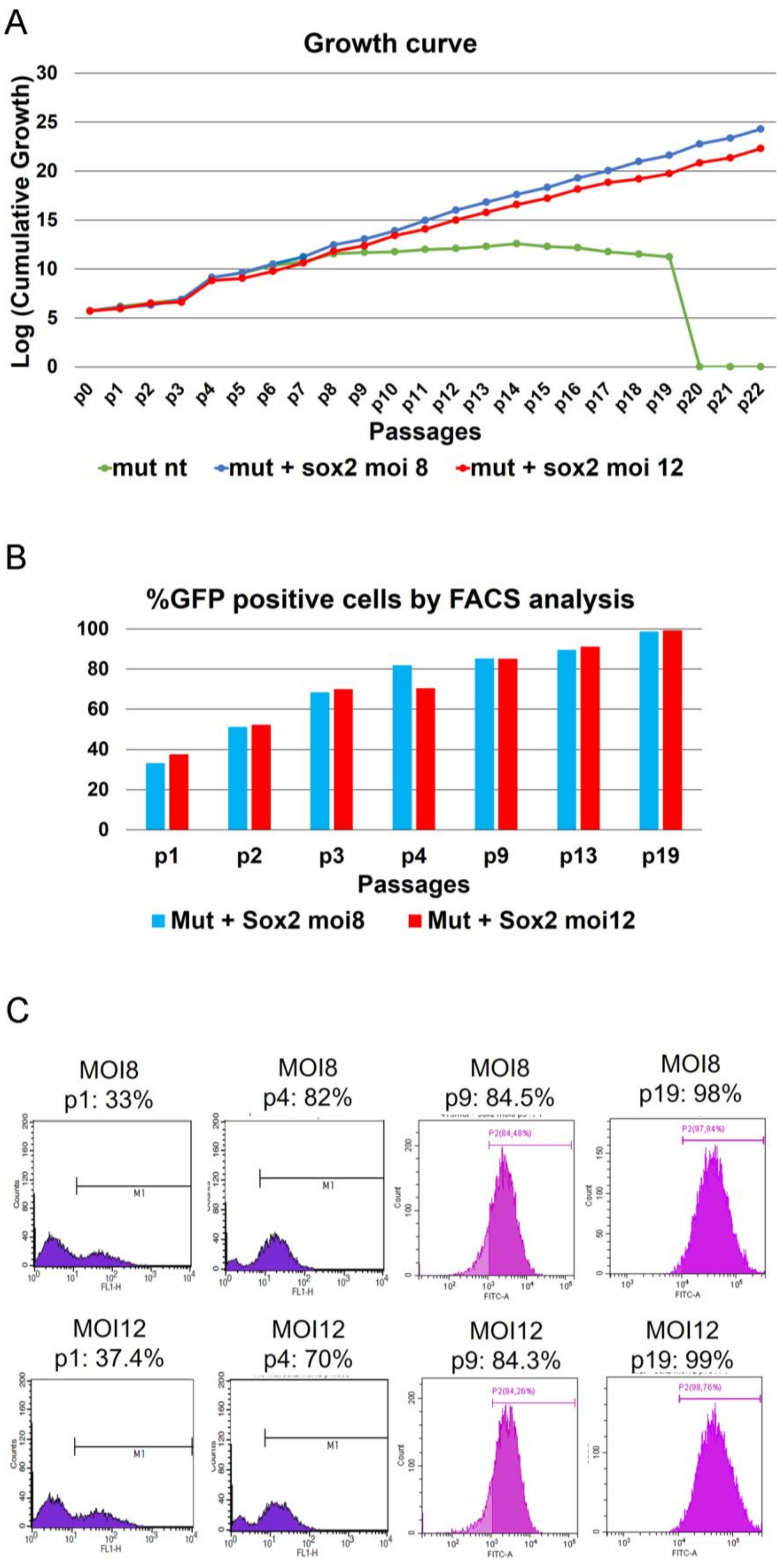

**Supplementary Fig. 4**

**Rescue of long-term proliferation of Sox2-deleted NSC by lentiviral Sox2 transduction**

**A,** Growth curve of Sox2-deleted (mut) cells, untransduced (green) or transduced with Sox2- and GFP-expressing lentivirus at MOI 8 (blue) or 12 (red).

**B,** Percentage of GFP-positive cells at different passages

**C,** Examples of FACS analyses at different passages

### Supplementary Figure 5

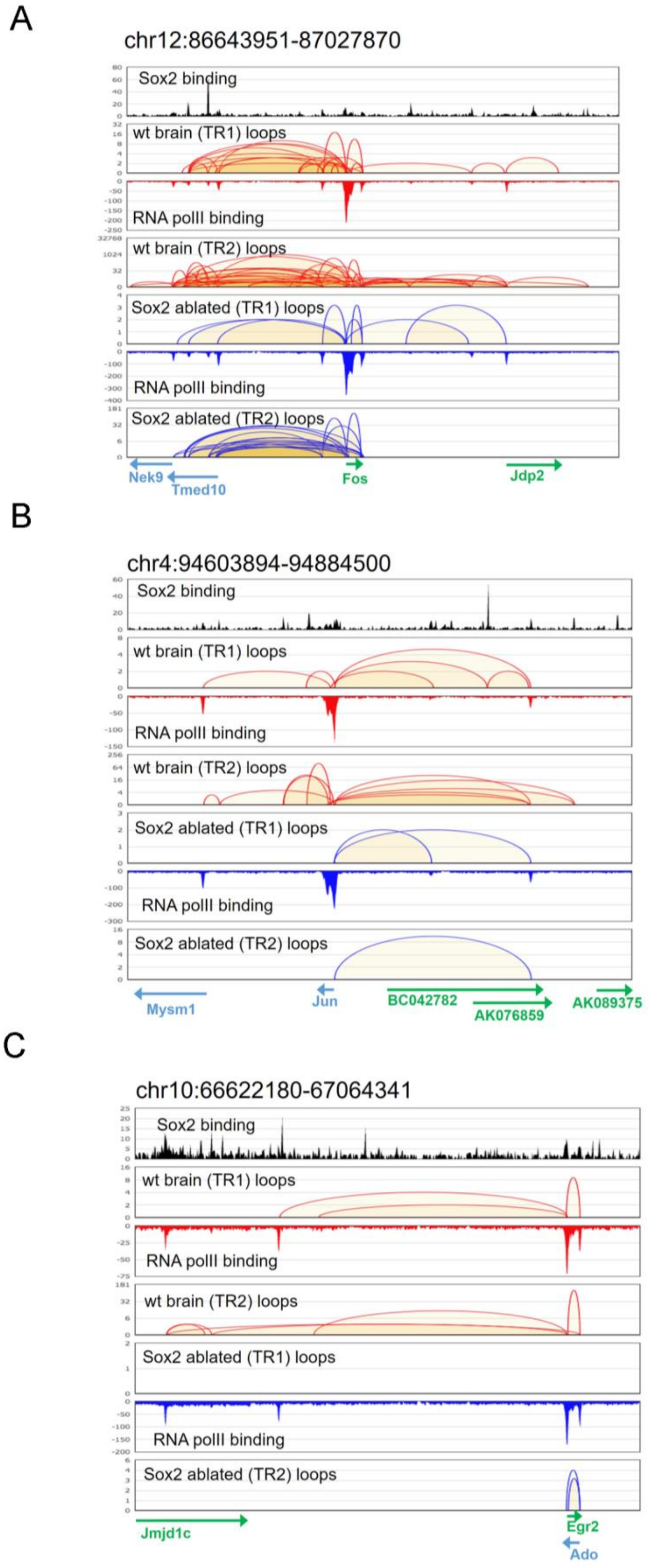

**Supplementary Figure 5**

**Long-range chromatin interactions detected by ChIA-PET on the Fos (A), Jun (B), and Egr2 (C) regions in wild type and Sox2-deleted NSC**

The Fos (A), Jun (B) and Egr2 (C) genes are indicated below each panel. Long-range interactions between the genes and distal regions are indicated by loops (wild type cells, red; Sox2-deleted cells, blue). Tracks with SOX2 and RNApolII binding peaks are shown. TR1 and TR2 refer to different ChIA-PET experiments. Note the occurrence of SOX2 binding peaks in correspondence of distant regions connected with each of the three genes. Data from [8].

**Supplementary Table 1**

| | Mutant bacterial clones | | | Mutant alleles in cellular clones |
|---|---|---|---|---|
| | 36h after puromycin selection | 8 days after puromycin selection | 20 days after puromycin selection | |
| Mut | 9/10 tot | 10/10 | 16/16 tot | 14/18 tot * |
| Mock | 0/3 tot | | | 0/8 |

* One of the Mut clones was wild type homozygous

**Supplementary Table 1**

**Number of mutant/total alleles of Fos following CRISPR/Cas9 mutagenesis**

DNA was extracted from cells treated with lentiviruses at 36 hours, 8 days, and 20 days after puromycin selection; the Fos gene region containing the sgRNA-targeted sequence was amplified by PCR, and cloned into a pGEM®-T Easy plasmid. Bacterial clones were individually grown, plasmid DNA was extracted, and the plasmid insert was sequenced. The number of clones carrying a mutated Fos region is reported, over the total number of sequenced clones. The number of mutant alleles was also determined in individual cellular clones obtained after long-term propagation. Mutant clones were not found in Mock-transduced controls.

**Supplementary Table 2**

| | Experiment 1 | | Experiment 2 | |
|---|---|---|---|---|
| | Mut | Mock | Mut | Mock |
| Clones collected from 96-well plate | **23** | **11** | **25** | **20** |
| Clones which died in 48-well plate | 6 | 0 | 9 | 0 |
| Clones that do not proliferate after seeding in 12-well plate | 10 | 3 | 8 | 4 |
| Clones that continue to proliferate (in 6-well plate or in flask) | **7** | **8** | **8** | **16** |

**Supplementary Table 2**

**Comparison between antiFos Cas9-treated and Mock-treated NSC, as to their ability to generate clones capable of long-term expansion**

Numbers of primary clones in two experiments (clones from 96-well plates) obtained for further replating for long-term growth analysis (first row). After replating, some of the clones continued to efficiently proliferate long-term in 6-well plates or in flasks, for at least two months (fourth row). Only these clones are scored as derived from long-term self-renewing NSC.

**Supplementary Table 3**

| sgRNA anti-Fos 5'-CCGCTGCAGTAGCGCCTCCCCGG-3' | | | |
|---|---|---|---|
| Potential off-target sequence | | Number of mismatches | Genomic coordinates |
| A | 5'-C**A**GCTG**T**AGTAGCGCCTC**T**CAGG-3' | 3 | Chr10: 109101673-109101695 |
| B | 5'-CCGC**A**GCAG**CC**GCGCCTCCCCGG-3' | 3 | Chr3: 21935959-21935981 |
| C | 5'-CCG**G**T**T**CAGTAGCGCCT**T**CCTGG-3' | 3 | Chr15: 79511971-79511993 |
| D | 5'-CCGCTGCAGT**G**GC**CG**CTCCCTGG-3' | 3 | Chr12: 76962036-76962058 |
| E | 5'-C**A**GCTGCAGTAGCGC**TG**CCCCGG-3' | 3 | Chr15: 102937211-102937233 |
| F | 5'-CC**T**CTGCAGTAG**GA**CCTCCCGGG-3' | 3 | Chr8: 120361641-120361663 |

**Supplementary Table 3**

**Potential CRISPR/Cas9 off-target sequences**

The potential CRISPR/Cas9 off-targets sequences (called A to F), with their genomic coordinates, carrying an appropriately located PAM sequence (highlighted in gray), and bearing similarity to the guideRNA used (sgRNA anti-Fos). All these sequences had 3 mismatches with the anti-Fos sgRNA.

**Supplementary Table 4**

| Potential off-target sequence | Total plasmids analyzed | Mutant plasmids | Mutations |
|---|---|---|---|
| A | 8 | 0 | |
| B | 10 | 0 | |
| C | 8 | 0 | |
| D | 21 | **1** | 5'-CCGCTGCAGTGGCCGCTC**T**CTGG-3' |
| E | 9 | 0 | |
| F | 7 | 0 | |

**Supplementary Table 4**

**Analysis of mutations in CRISPR/Cas9 off-targets regions**

The potential CRISPR/Cas9 off-targets regions A to F were PCR-amplified, cloned into plasmids and individually sequenced (the total number of plasmids analyzed is shown). Only one mutation in region D (1 out of 21 different plasmids sequenced) was identified.

**Supplementary Table 5**

| N° nucleotides | Insertions | Deletions |
|---|---|---|
| 1 | 5 | 2 |
| 2 | 5 | 1 |
| 3 | 1 | 2 |
| 4 | | 1 |
| 5 | 1 | 3 |
| 6 | | |
| 7 | | |
| 8 | | 1 |
| 9 | | 3 |
| 10 | | 1 |
| 11 | | |
| 12 | | 1 |
| 13 | | |
| 14 | | |
| 15 | | |
| 16 | | |
| 17 | | 1 |
| 18 | | |
| 19 | | 1 |
| 20 | | |
| 21 | 1 | |
| 22 | | |
| 23 | | |
| 24 | | 2 |
| 25 | | |
| 26 | | |
| 27 | | |
| 28 | | |
| 29 | | |
| 30 | | |
| 31 | | 1 |
| 32 | | |
| 33 | | |
| 34 | | |
| Total | 13 | 20 |
| Total indels | 33 | |
| Total alleles | 35* | |

*two of the sequenced alleles maintain the correct reading frame, but are extensively mutagenized around the sgRNA-complementary region.

**Supplementary Table 5**

**Indel mutations induced by CRISPR/Cas9-antiFos sgRNA in the Fos gene**
